# Re-enacting steps supports human path integration consistent with motor-corrected grid cell drift

**DOI:** 10.1101/2025.05.18.654711

**Authors:** Volker Reisner, Leonard König, Viktor Studenyak, Marcia Bécu, Andrej Bicanski, Christian F. Doeller

## Abstract

Efficient navigation, especially in the absence of vision, requires path integration — the continuous updating of spatial position from self-motion cues. However, path integration is prone to cumulative error, which can be amplified when body-derived information is inconsistent between encoding and retrieval paths. Drawing on evidence from sensorimotor reactivation during memory retrieval, we hypothesized that re-enacting encoding-related movement patterns could serve as a body-derived mechanism to counteract such errors. In a novel virtual reality task with motion tracking, participants learned unique, irregular step sequences linked to specific target distances during an encoding phase. They later reproduced these distances in complete darkness under three conditions: self-paced free retrieval, retrieval with encoding-congruent movements, and retrieval with encoding-incongruent, regular gait. During free retrieval, participants naturally re-enacted encoding-related movements associated with improved distance reproduction. As predicted, distance estimation was significantly more accurate during congruent retrieval than during incongruent retrieval. These behavioral findings are consistent with a neural network model in which retrieved encoding-related motor patterns correct path integration errors in grid cells. Together, these results provide converging behavioral and computational evidence that body-derived, encoding-related motor patterns can enhance distance estimation, possibly by filtering grid cell error accumulation, offering new insights into embodied mechanisms of path integration.

## Introduction

A fundamental problem in spatial navigation is to return to a start location after reaching out. To achieve this, humans and other animals use *path integration* (PI), which continuously updates their location based on self-motion information (Etienne & Jeffrey, 2004). In the brain, this process is likely supported by grid cells in the medial entorhinal cortex (MEC; Moser, Rowland & Moser, 2015; McNaughton et al., 2006), which exhibit a highly regular, lattice-like firing pattern (Hafting et al., 2005). However, due to the recursive nature of PI, grid cells accumulate error over time, leading to drift in their firing pattern (Hardcastle et al., 2015). Similarly, human PI error steadily increases with distance (e.g. Durgin et al., 2009; Lappe, Jenkin & Harris, 2007). While unchecked error accumulation can cause an eventual collapse of the grid code (Burak & Fiete, 2009), stable environmental cues can provide effective compensation (Hardcastle et al., 2015).

But how is PI maintained in the absence of environmental cues, as in darkness? In humans, non-visual PI relies primarily on the vestibular and haptic systems (idiothetic cues) derived from self-motion (Mittelstaedt & Mittelstaedt, 2001). Research has shown that blindfolded participants can reliably reproduce distances up to 100 meters when the gait pattern used on the return matches the outbound gait (Bigel & Ellard, 2000; Durgin et al., 2009; Glasauer, Amorim & Berthoz, 1994; Klatzky et al., 1990; Loomis et al., 1999; Mittelstaedt & Mittelstaedt, 2001; Schwartz, 1999). In contrast, errors increase when the type of gait is changed across paths, e.g. from walking to galloping (Abdolvahab et al., 2015; Chrastil & Warren, 2014; Harrison, 2020; Harrison & Davis, 2023; Harrison et al., 2021; Turvey et al., 2009, 2012), suggesting that distance reproduction heavily relies on encoding-related body-derived signals.

From a memory perspective, the outbound path represents the encoding phase, during which information is gathered to estimate the distance traveled, whereas the return path mirrors retrieval, drawing on perceptual and motor information of the original experience (cf. Ianì, 2019). Consistent with this view, a wide range of neuroimaging studies showing overlapping cortical activation during encoding and retrieval of sensory and motor information (Danker & Anderson, 2010; Kent & Lamberts, 2008; Masumoto et al., 2006; Nilsson et al., 2000; Nyberg et al., 2001) as well as reinstatement in premotor cortex coupled with the hippocampus (Meyer et al., 2024). The hippocampus has also been linked to the reactivation of memories associated with motor sequences (Albouy et al., 2015; Döhring et al., 2017; Jacobacci et al., 2022). Behaviorally, sensorimotor reactivation predicts that re-enacting encoding-related sensorimotor states facilitates retrieval, while those interfering with the encoded states can hinder it. Re-enactment effects have been previously observed for various domains such as body posture (e.g. Dijkstra, Kaschak & Zwaan, 2007; Morse et al., 2015), gestures (e.g. Iverson & Goldin-Meadow, 1998; Ping, Goldin-Meadow & Beilock, 2014; Morsella & Krauss, 2004), and eye movements (e.g. Wynn, Shen & Ryan, 2019).

An interesting scenario in which PI may heavily rely on sensorimotor reactivation arises when locomotion patterns themselves are explicitly encoded. Here, estimating distance may be fundamentally different from perceptually-driven odometry. That is, when a sequence of steps forms a complex spatio-temporal pattern, the necessity of higher-order cognitive processes is pronounced, where spatial memory reflects the relative location of foot placements and motor sequence memory ensures correct step execution with the appropriate leg. At the neural level, spatial memory is closely linked to neural codes in the entorhinal-hippocampal circuit, including MEC neurons, which play an important role in the representation of linear distance in total darkness (Campbell et al., 2021; Jacob et al., 2017). In contrast, motor sequence memory is primarily supported by the motor network (cf. Magill, 2011; Schmidt & Lee, 2011) and the hippocampus which represents information about the learned temporal order of sequential motor actions (e.g. Albouy et al., 2008; Dolfen et al., 2024; Yewbrey & Kornysheva, 2024). Therefore, if complex, irregular movement sequences (motor cortex) are explicitly encoded with target distances (MEC) within memory (hippocampus), then the re-enactment of those encoding-related motor patterns during retrieval may provide an internal corrective signal, even in the absence of environmental cues. Thus, we predicted reduced distance reproduction errors when motor patterns mimic encoded movements compared to different movements.

In the present study, we explored the role of behavioral re-enactment in estimating the distance travelled via discrete, irregular movement patterns, and simulated how specific memorized motor patterns could interact with PI. Using a novel distance reproduction task, participants learned to associate unique motor sequences with target distances of varying length before reproducing them in total darkness with movement patterns either congruent or incongruent to encoding. We found that distance reproduction was strongly influenced by the degree of re-enactment and disrupted by movement changes between encoding and retrieval. Our neurocomputational model suggests how motor-driven error correction of grid cell-based spatial representations can account for our findings.

## Results

We used immersive virtual reality (VR) and motion capture (MoCap) technology to investigate the role of encoding-related movement sequences in non-visual path integration (PI). 29 participants (12 women, 14 men, 3 non-binary; age: Mean = 26.86 years; SD = 4.45 years, range = 19-35 years) performed a novel distance reproduction task that comprised a training and a test phase (Fig. 1a). During training (Fig. 1b, left), participants learned target locations from different distances (short, medium, long) from a fixed start location, each associated with unique irregular step patterns along a virtual 1-D linear track (Fig. 1c). During subsequent test (Fig. 1b, right), participants were asked to return to the target locations in complete darkness by (1) walking freely without any movement-related instruction (‘free retrieval’), (2) reproducing encoding-congruent step patterns (‘congruent retrieval’), and (3) executing regular, encoding-incongruent steps (‘incongruent retrieval’).

**Figure 1.**
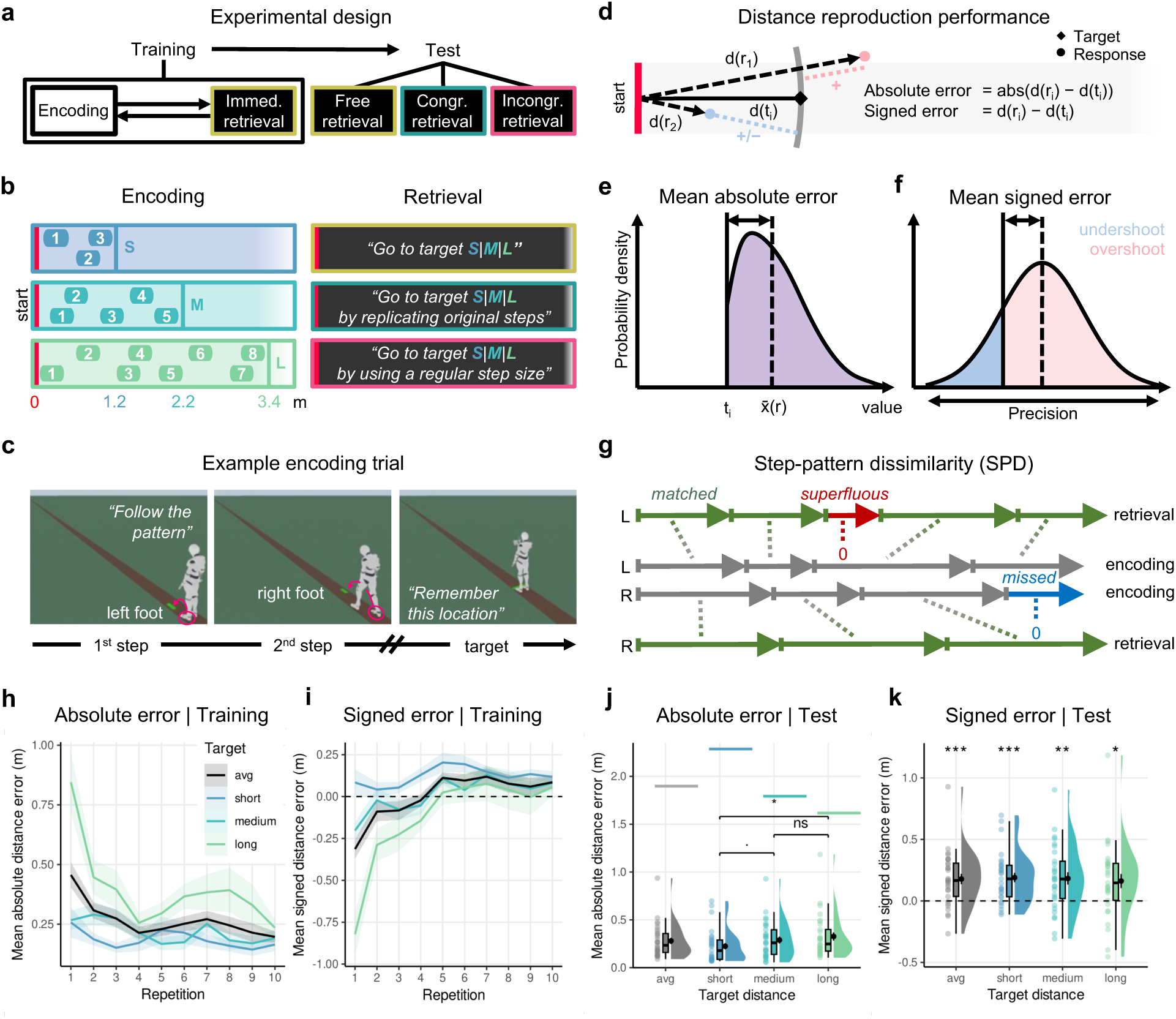
Task design, measures and distance reproduction performance. **a-c:** Task design. **a:** Participants performed a distance reproduction task consisting of a training phase with alternating encoding and immediate retrieval trials, as well as a test phase with three retrieval conditions. **b:** During training (left), participants encoded target locations at varying distances (S = short, M = medium, L = long) from a fixed start location by following unique irregular step patterns along a virtual linear track, with no proximal or distal cues. During the test phase (right), participants returned to the targets in complete darkness under three retrieval conditions: (1) free retrieval, i.e. walking without specific movement instructions, (2) congruent retrieval, i.e. walking that mimics the original step pattern, and (3) incongruent retrieval, i.e. walking with a regular step size. Task instructions during free retrieval were the same as during immediate retrieval during training **c:** Example encoding trial. Alternating left and right footholds dynamically appeared along the linear track. As soon as a step was automatically detected, it disappeared and the next step was introduced until the target location (a colored tag on the floor) was reached. The participant’s body was virtually represented as an individually scaled avatar. **d-f:** Measures (dependent variables). **d:** On each trial, distance reproduction performance was measured as the difference between start-to-target (diamond) and start-to-response (circle) distances. Absolute error reflects unsigned deviations, while signed error indicates deviation direction (red/blue). **e**: Across trials, the mean absolute distance error *x̅*(*r*) reflects the sum of both positive and negative mean deviations (purple) from a target ti (gray line). **f:** The mean signed distance error is positive when targets were systematically overshot (red) and negative when they were undershot (blue), with its variation indicating the precision. **g**: The *step pattern dissimilarity* (SPD) quantifies how closely leg movements during retrieval resembled those during encoding: Differences between encoded and retrieved step lengths matched by Dynamic Time Warping (dotted lines) are summed and non-matching steps (superfluous and missed) are added to the score. **h-k:** Distance reproduction performance during training (immediate retrieval) and test (free retrieval) for short (blue), medium (cyan), and long (green) target distances, as well as the average across targets (black). **h:** Mean absolute distance error during training decreased with repetition, particularly in long distance trials. Colored lines reflect the mean ± SEM (ribbon) per condition. **i:** Mean signed distance error during training was negative during the first half of trials and positive during the second half. **j:** Mean absolute distance error during test with target-specific chance levels (colored horizontal lines). Mean error was below chance and higher for long distance vs short distance. Violin plots depict the density distribution, boxplots the median and quartiles, black dots with error bars the means ± SEM, and colored dots individual participants per condition. **k:** Mean signed distance error during test with target-specific chance levels (colored horizontal lines). Positive mean error indicated systematic overshooting across target distances. ᐧ *P* < .10, * P < .05.

### Distance reproduction performance

To assess participants’ ability to reproduce previously experienced distances in complete darkness, we examined (immediate/free) retrieval trials during training and test, both in which no movement-related instruction was given (Fig. 1a-b). For each trial, we measured the degree of overshooting or undershooting of target distances and determined the absolute and signed distance error for all target distances and on average (Fig. 1d-f).

During training, all participants completed a total of 10 repetitions per target distance. A bootstrapped 2-way repeated measures ANOVA on absolute distance errors revealed significant main effects of repetition (*F*_3.7,103.7_ = 8.41, *P* < 0.0001, η^2^_*p*_= 0.23, 95% CI = 0.14 to 0.39) and target distance (*F*_1.2,32.2_ = 24.2, *P* < 0.0001, η^2^_*p*_= 0.46, 95% CI = 0.35 to 0.76), as well as their interaction (*F*_4.8,135.4_ = 4.03, *P* < 0.0001, η^2^_*p*_= 0.13, 95% CI = 0.09 to 0.25; Fig. 1h). On average, participants reduced their error by 0.26 m (SD = 0.28 m) from the first to the last repetition (bootstrapped paired *t*-test: *t*_28_ = 5.03, *P*_adj_ < 0.001, *d* = 0.93, 95% CI = 0.7 to 1.3, Bonferroni-corrected for 44 comparisons) with long distance accumulated more error than short distance (bootstrapped paired *t*-test: *t*_28_ = -5.30, *P*_adj_ < 0.001, *d* = -0.98, 95% CI = -1.95 to -0.76, Bonferroni-corrected for 3 comparisons). Despite fluctuations over time, distance reproduction of long distance consistently showed the highest improvement between the first and last repetition (short: M = -0.09, SD = 0.39; medium: M = -0.08, SD = 0.29; long: M = -0.61, SD = 0.63). Notably, mean signed distance errors shifted from negative to positive (Fig. 1i), indicating undershooting during the first half of learning (bootstrapped 1-sample *t*-test against 0: *t*_28_ = -3.15, *P*_adj_ < 0.01, *d* = -0.59, 95% CI = -0.95 to -0.29, Bonferroni-corrected for 2 comparisons) and overshooting during the second half of trials (*t*_28_ = 2.77, *P*_adj_ < 0.01, *d* = 0.51, 95% CI = 0.14 to 1.06, Bonferroni-corrected for 2 comparisons).

Following training, participants’ distance reproduction performance was probed in free retrieval trials during the test phase (where no feedback was provided and movement was self-paced). For each target distance, we tested participants’ absolute distance errors against chance levels derived from 1,000 artificial participants responding at random locations along the linear track, each preserving the original trial structure (see “Methods” under “Data processing and analysis”; Supplementary Fig. S2a). Bootstrapped 1-sample *t*-tests verified that errors fell far below the chance level for each target distance (all *P*_adj_ ≤ 0.0001, Bonferroni-corrected for 3 comparisons; Fig. 1j), as well as on average (*t*_28_ = -49.75, *P* < 0.0001, *d* = -9.24, 95% CI: -16.47 to -6.41). Similarly to training, long distance accumulated more error compared to short distance (bootstrapped paired *t*-test: *t*_28_ = -3.02, *P*_adj_ = 0.014, *d* = -0.56, 95% CI = -1.02 to -0.2, Bonferroni-corrected for 3 comparisons), whereas the mean signed of error was consistently positive for all target distances (bootstrapped 1-sample *t*-test against 0: all *P*_adj_’s ≤ 0.02) and on average (*t*_28_ = 4.23, *P* < 0.001, *d* = 0.79, 95% CI: 0.48 to 1.31), indicating systematic overshooting of target distances (Fig. 1k), consistent with previous studies including target distances less than approximately 10 meters (e.g. Klatzky et al., 1990; Loomis et al., 1993; Marlinsky, 1999; Schwartz, 1999; Turvey et al., 2009). While participants’ reproduced distance was roughly a linear function of target distance (Supplementary Fig. S3a), its precision was fairly constant with coefficients of variance in the logarithmic domain being similar across target distances (bootstrapped 1-way repeated measures ANOVA: *F*_1.9,52.8_ = 1.22, *P* = 0.31, η^2^_*p*_= 0.04, 95% CI = 0.003 to 0.23; Supplementary Fig. S3b). This resonates with Weber’s law, which states that the just-noticeable difference in a physical property is a fixed proportion of its magnitude (Foster, 1923) and aligns with previous observations focusing on regular walking (Durgin et al., 2009). Overall, these data show that participants learned to reproduce all three target distances properly.

### Uninstructed re-enactment of encoding-related step patterns facilitates distance reproduction

What role do step patterns play in the memory-guided reproduction of distances when movement is fully unrestricted? To explore this, we compared various movement characteristics during encoding with those during free retrieval (Fig. 2). Figure 2a shows example step patterns, one for each target distance.

**Figure 2.**
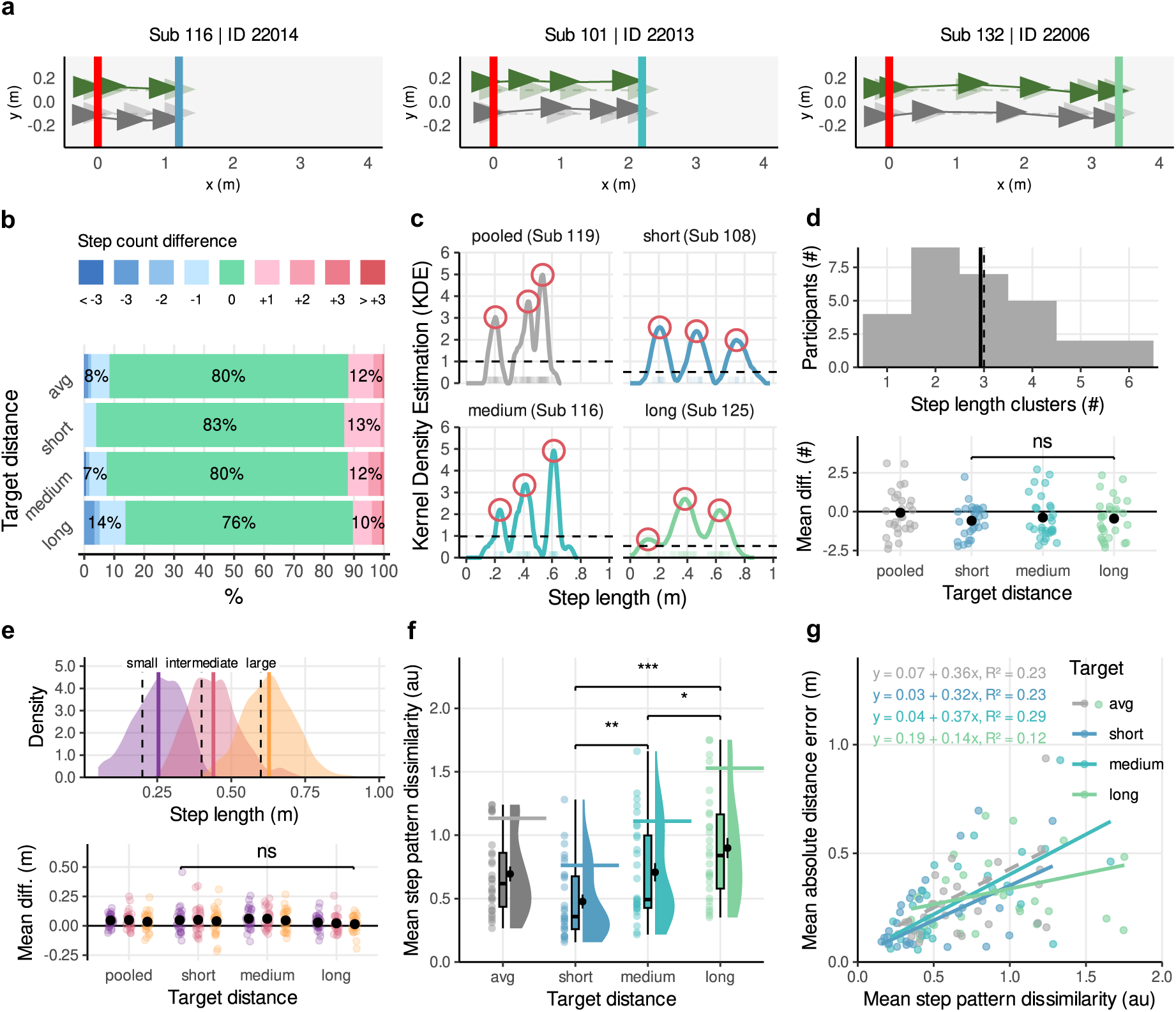
Movement characteristics during free retrieval. **a:** Step sequences of 3 example participants for a short (left), medium (middle) and long (right) target distance trial. Participants started from a fixed location (red line at 0 m) and executed left (dark green) and right (gray) steps until they stopped (response). Transparent triangles represent encoded steps towards the target location (blue, cyan, light green lines). **b-g:** Movement parameter for each target distance (short, medium, long, and averaged/pooled). **b:** Step count difference between encoding and retrieval (percentage). On average, in 80% of trials participants matched the number of performed steps between encoding and retrieval. **c:** Example Kernel Density Estimation of 4 participants. Red circles marking detected cluster peaks and the dashed line denotes a relative threshold of 20% of the maximum magnitude. **d:** *Top:* Frequency of participants as a function of step length clusters (bars) identified within pooled step length KDEs. The mean number of clusters (solid line) aligned with the number of encoded step lengths (dashed line). *Bottom:* The mean difference from 3 (true number) was comparable across target distances. Black dots with error bars depict the means ± SEM, and colored dots individual participants per condition. **e:** *Top:* Density of pooled step lengths labeled as small (purple), intermediate (red) and large (orange). The mean of each labelled step length (solid lines) was slightly greater than true step lengths (dashed lines). *Bottom:* The differences between labeled and true step lengths were comparable across cluster labels and target distances. **f:** Mean step pattern dissimilarity with target-specific chance levels (colored lines). Step patterns dissimilarity was below chance level and decreased with target distance. Violin plots depict the density distribution, boxplots the median and quartiles. **g:** Relationship between mean step pattern dissimilarity and mean absolute distance error. Kendall-Theil regressions revealed a positive linear relationship, indicating the more participants re-enacted encoding-related steps, the better their distance reproduction. * *P* < .05, ** *P* < .01, *** *P* < .001.

First, we calculated the difference in the number of steps between encoding and retrieval for each participant and tested whether these differences were deviating from zero. On average, participants performed the same number of steps in 80% of trials (bootstrapped 1-sample *t*-test on step count difference against 0: *t*_28_ = 0.48, *P* = 0.65, *d* = 0.09, 95% CI = -0.23 to 0.568; Fig 2b) with no significant differences between target distances (bootstrapped 1-way repeated measures ANOVA: *F*_1.7,46.7_ = 1.68, *P* = 0.2, η^2^_*p*_= 0.06, 95% CI = 0.003 to 0.24). Next, we identified the number of clusters within the Kernel density estimation (KDE) of step lengths for each participant (Fig. 2c) and compared these to the ground truth of *N* = 3 encoded step lengths. Considering all the data, we found a mean number of approximately 3 clusters (M = 2.93, SD = 1.41), consistent with the true number of encoded step lengths (bootstrapped 1-sample *t*-test against 3: *t*_28_ = -0.26, *P* = 0.8, *d* = -0.05, 95% CI = -0.47 to 0.3; Fig. 2d, top), with no differences across target distances (bootstrapped 1-way repeated measures ANOVA: *F*_1.87,52.27_ = 0.2, *P* = 0.82, η^2^_*p*_= 0.007, 95% CI = 0.001 to 0.14; Fig. 2d, bottom). According to this number, we further assigned all step lengths to small, intermediate, and large step length classes using the Jenks Natural Breaks Classification (Jenks, 1967) and calculated the deviation of each class’ mean from the corresponding encoded step length. Among all step lengths, we found the highest positive deviation for small steps (bootstrapped 1-sample *t*-test against 0: *t*_28_ = 4.49, *P*_adj_ < 0.001, *d* = 0.83, 95% CI = 0.48 to 1.34, Bonferroni-corrected for 3 comparisons) and the smallest deviation for large steps (*t*_28_ = 2.38, *P*_adj_ = 0.02, *d* = 0.44, 95% CI = 0.09 to 0.88, Bonferroni-corrected for 3 comparisons; Fig. 2e, top). However, a bootstrapped 2-way repeated measures ANOVA revealed comparable deviation across step length classes (*F*_1.8,49.4_ = 0.96, *P* = 0.39, η^2^_*p*_= 0.03, 95% CI = 0.003 to 0.27), target distances (*F*_2.2,62.8_ = 2.43, *P* = 0.08, η^2^_*p*_= 0.08, 95% CI = 0.02 to 0.23), and no interaction (*F*_2.7,74.4_ = 0.10, *P* = 0.99, η^2^_*p*_= 0.003, 95% CI = 0.005 to 0.13; Fig. 2e, bottom). Ultimately, we calculated the step pattern dissimilarity (SPD) that quantifies the mismatch of step patterns between encoding and retrieval by taking into account the order of executed steps (Fig. 1g). As a benchmark, we established a target-specific chance level by simulating SPD values based on random step patterns falling between the start and target locations for 1,000 artificial participants, each maintaining the empirical trial structure (see “Method” under “Data processing and analysis”; Supplementary Fig. S2b). Bootstrapped 1-sample *t*-tests confirmed that the participants’ SPD were significantly below chance for all target distances (all *P*_adj_’s ≤ 0.001, Bonferroni-corrected for 3 comparisons), and on average (*t*_28_ = -7.48, *P* < 0.0001, *d* = -1.39, 95% CI = -2.19 to -0.96; Fig. 2f). As expected, SPD values increased as a function of target distance (bootstrapped 1-way repeated measures ANOVA: *F*_1.7,46.7_ = 22.73, *P* < 0.0001, η^2^_*p*_= 0.45, 95% CI = 0.34 to 0.59), reflecting the cumulative nature of step deviations. Importantly, Kendall-Theil regressions demonstrated that SPD values were robust predictors of absolute distance error across all target distances (all slope *P*’s ≤ 0.002, Bonferroni-corrected for 3 comparisons; all *R*^2^_*E*_ ≥ 0.12) and on average (β_1_ = 0.36, 95% CI = 0.21 to 0.39, *V* = 419, *P* < 0.0001, *R*^2^_*E*_ = 0.23; β_0_ = 0.07, 95% CI = 0.04 to 0.11, *V* = 365, *P* < 0.001; Fig 2g). This indicates that the accuracy by which participants reproduced distances improved as the similarity in step patterns between encoding and retrieval increased. In sum, our movement analysis suggests that participants intuitively re-enacted encoding-related step patterns, which in turn facilitated distance reproduction.

### Suppressing encoding-related re-enactment disrupts distance reproduction

To directly test how the (mis)alignment of step patterns between encoding and retrieval affects distance reproduction, we compared retrieval trials where participants were instructed to either replicate encoding-congruent steps or use regular, encoding-incongruent steps. Exemplary step patterns for congruent and incongruent retrieval trials are shown in Figure 3a.

**Figure 3.**
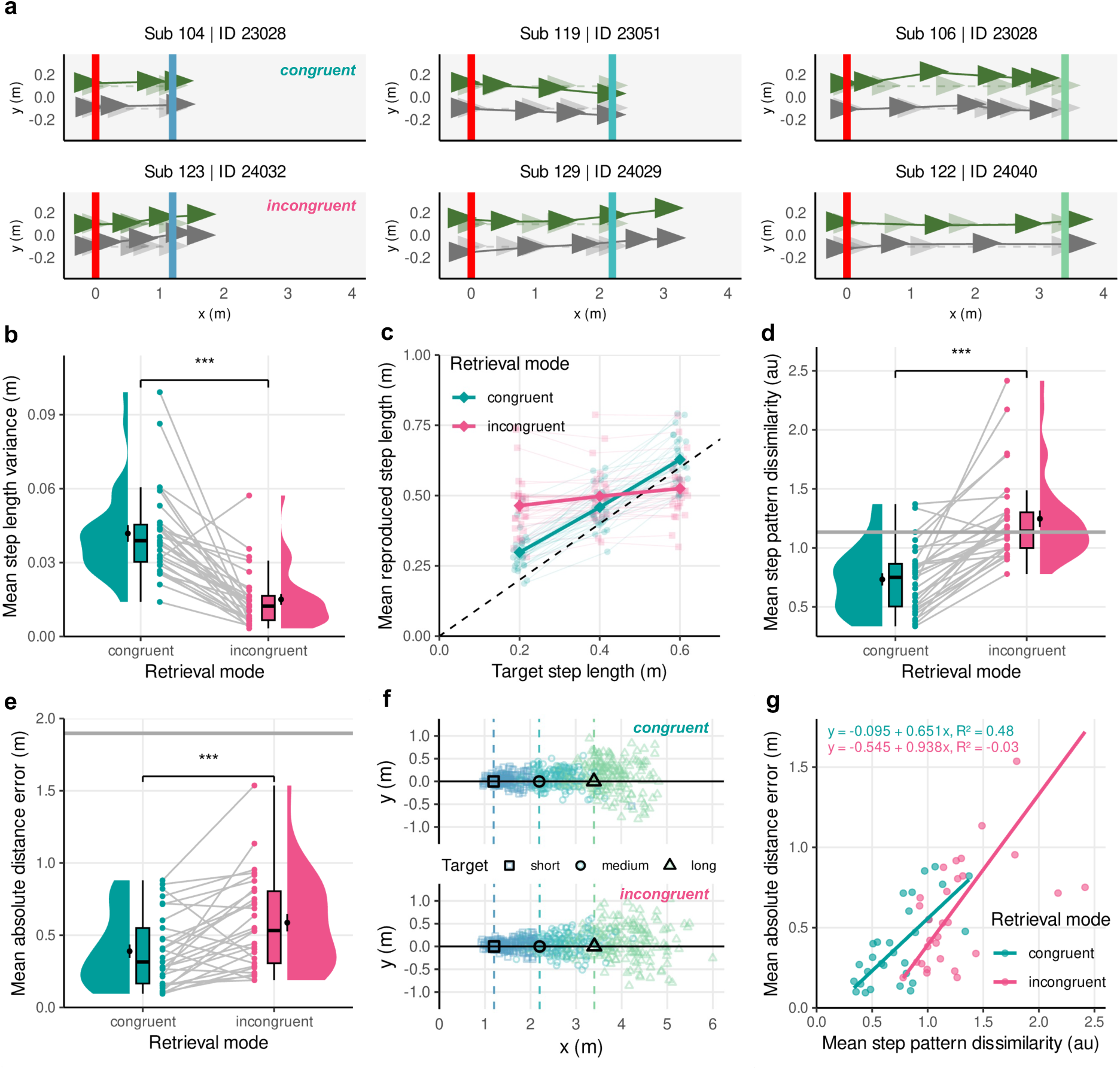
Movement and distance reproduction during congruent vs incongruent retrieval. **a:** Step sequences of 3 example participants for a short (left), medium (middle) and long (right) target distance trial during congruent (top) and incongruent retrieval (bottom). Participants started from a fixed location (red line at 0 m) and executed left (dark green) and right (gray) steps until they stopped (response). Transparent triangles represent encoded steps towards the target location (blue, cyan, light green lines). **b-d:** Movement parameters for congruent (green) vs incongruent (magenta). **b:** Step length variance. Higher variation in step length during congruent (irregular step patterns) vs incongruent (regular step patterns). Violin plots depict the density distribution, boxplots the median and quartiles, black dots with error bars the means ± SEM, and colored dots individual participants per condition. **c:** Mean reproduced step length as a function of target step length aligned by Dynamic Time Warping (DTW) that minimizes the distance between the two step sequences. Reproduced step lengths matched target step lengths better during congruent than incongruent retrieval. Note: One participant was excluded for whom no incongruent step was classified as small (< 0.05 m). Bold dots with error bars the means ± SEM, and colored dots individual participants (jittered) per condition. Dotted line represents a perfect linear relationship. **d:** Mean step pattern dissimilarity. Dissimilarity was higher during incongruent than congruent retrieval, with congruent values falling below chance (gray line), unlike incongruent. **e-g:** Distance reproduction performance for congruent (green) vs incongruent (magenta). **e:** Mean absolute distance error. Participants showed greater errors during incongruent retrieval than congruent retrieval, yet performed above chance (gray line) in both conditions. **f:** Raw responses (color-coded dots) for congruent (top) and incongruent (bottom) retrieval, separated by target distance (symbols; dashed lines) along the linear track. **g:** Relationship between mean step pattern dissimilarity and absolute distance error during congruent vs incongruent retrieval. Kendall-Theil regressions revealed strong explanatory power of SPD values on distance reproduction for congruent, but not incongruent retrieval. *** *P* < .001.

In order to verify that participants were following movement-related instructions, we first compared movement characteristics across conditions (Fig. 3b-d). For this, we calculated the variance within sequences in the length of executed steps for each participant. A bootstrapped paired *t*-test revealed a significant difference with greater step length variance during congruent than incongruent retrieval (*t*_28_ = 6.9, *P* < 0.0001, *d* = 1.24, 95% CI = 0.88 to 2.01; Fig. 3b). Next, we took a closer look on how participants produced step lengths with respect to the encoded step lengths. To this end, we determined the closest match between produced encoded step lengths using Dynamic Time Warping (DTW) and calculated the mean for each participant. A bootstrapped 2-way repeated measures ANOVA yielded a significant interaction between the encoded step length and retrieval mode (*F*_1.5,41.4_ = 130.32, *P* < 0.0001, η^2^_*p*_= 0.83, 95% CI = 0.77 to 0.90), suggesting that reproduced step lengths were more closely aligned with encoded step lengths during congruent retrieval compared to incongruent retrieval (Fig. 3c). As a result of this match, participants exhibited lower levels of step pattern dissimilarity (SPD) between encoding and congruent retrieval than between encoding and incongruent retrieval (bootstrapped paired *t*-test: *t*_28_ = -7.4, *P* < 0.0001, *d* = -1.37, 95% CI = -2.18 to -1.03; Fig. 3d). Specifically, SPD values for congruent retrieval deviated significantly from chance (bootstrapped 1-sample *t*-test against chance: *t*_28_ = -7.67, *P*_adj_ < 0.001, *d* = -1.4, 95% CI = -2.26 to -0.96, Bonferroni-corrected for 2 comparisons) while those for incongruent retrieval did not (*t*_28_ = 1.62, *P*_adj_ = 0.27, *d* = 0.30, 95% CI = -0.05 to 0.57, Bonferroni-corrected for 2 comparisons). These results indicate that participants followed instructions by attempting to replicate encoding-related step patterns during congruent retrieval and using a more regular step length during incongruent retrieval.

We hypothesized that adopting a step pattern that differs from that used during encoding would impair distance reproduction performance. To test this, we compared absolute distance errors across retrieval modes using a bootstrapped paired *t*-test. Error levels were significantly higher during incongruent retrieval than during congruent retrieval (Absolute mean difference: M = 0.2 m, SD = 0.26 m; *t*_28_ = -3.81, *P* < 0.001, *d* = -0.71, 95% CI = - 1.22 to -0.35; Fig. 3e), which is apparent even in the raw responses (Fig. 3f). Notably, errors in both conditions fell significantly below the chance level (bootstrapped 1-sample *t*-tests for congruent: *t*_28_ = -31.14, *P*_adj_ < 0.001, *d* = -5.78, 95% CI = -8.16 to -4.66; incongruent: *t*_28_ = -15.94, *P*_adj_ < 0.001, *d* = -2.96, 95% CI = -5.98 to -2.00; Bonferroni-corrected for 2 comparisons), suggesting that even during incongruent retrieval, participants did not perform at random. Finally, we estimated the predictive power of SPD values on absolute distance errors for each retrieval mode using Kendall-Theil regressions (Fig 3g). We found that SPD values explained 48% of variance in distance reproduction performance when participants were walking congruent to encoding (β_1_ = 0.65, 95% CI = 0.49 to 0.76, *V* = 430, *P* < 0.0001, *R*^2^_*E*_ = 0.48; β_0_ = - 0.09, 95% CI = -0.13 to 0.0001, *V* = 127, *P* = 0.05). In contrast, the same regression model for the incongruent walking condition resulted in a negative coefficient of determination, suggesting that SPD values explained even less variance than the null model (β_1_ = 0.94, 95% CI = 0.57 to 1.15, *V* = 415, *P* < 0.0001, *R*^2^_*E*_ = -0.03; β_0_ = -0.55, 95% CI =-0.73 to -0.13, *V* = 88, *P* < 0.01). In summary, we demonstrated that suppressing the re-enactment of encoding-related step patterns by imposing regular gait diminished distance reproduction performance.

### Modelling error-correction for grid cells via stored motor patterns

To demonstrate how re-enacting encoding-related step patterns benefits non-visual distance reproduction, we developed a neurocomputational model that simulates the interaction between motor and memory systems. Specifically, we propose that the reactivation of motor activity can counteract error accumulation in a population of drifting grid cells that code for individual step placements. To directly compare the encoding and retrieval capabilities of our model with those of human participants, we simulated independent instances of the model, each corresponding to a simulated participant.

In the encoding phase (Fig. 4a, left, top), the model stores an association of performed motor patterns with corresponding spatial step locations represented by a grid cell population vector (PoV) in a modern Hopfield network (Ramsauer et al., 2021). Motor patterns were defined as unique tuples that capture the current and previous steps as well as their respective leg sides. Each of these tuples was assigned to a neuronal population, which represents a snapshot of activity in motor cortex. Step locations, on the other hand, were modeled as a grid cell PoV, constructed from a total of 252 neurons, by following the approach described in Bicanski and Burgess (2018). Given the functional role of modular organized grid cell activity for extracting distance information (Bush et al., 2015), the population of grid cells was modeled with 7 modules each containing 36 neurons, exhibiting varying frequencies, scaling, and orientations across modules and uniformly distributed spatial phases within each. To mimic the behavioral experiment (Fig. 1b), each simulated agent underwent a pseudo-random walk with varying target lengths.

**Figure 4.**
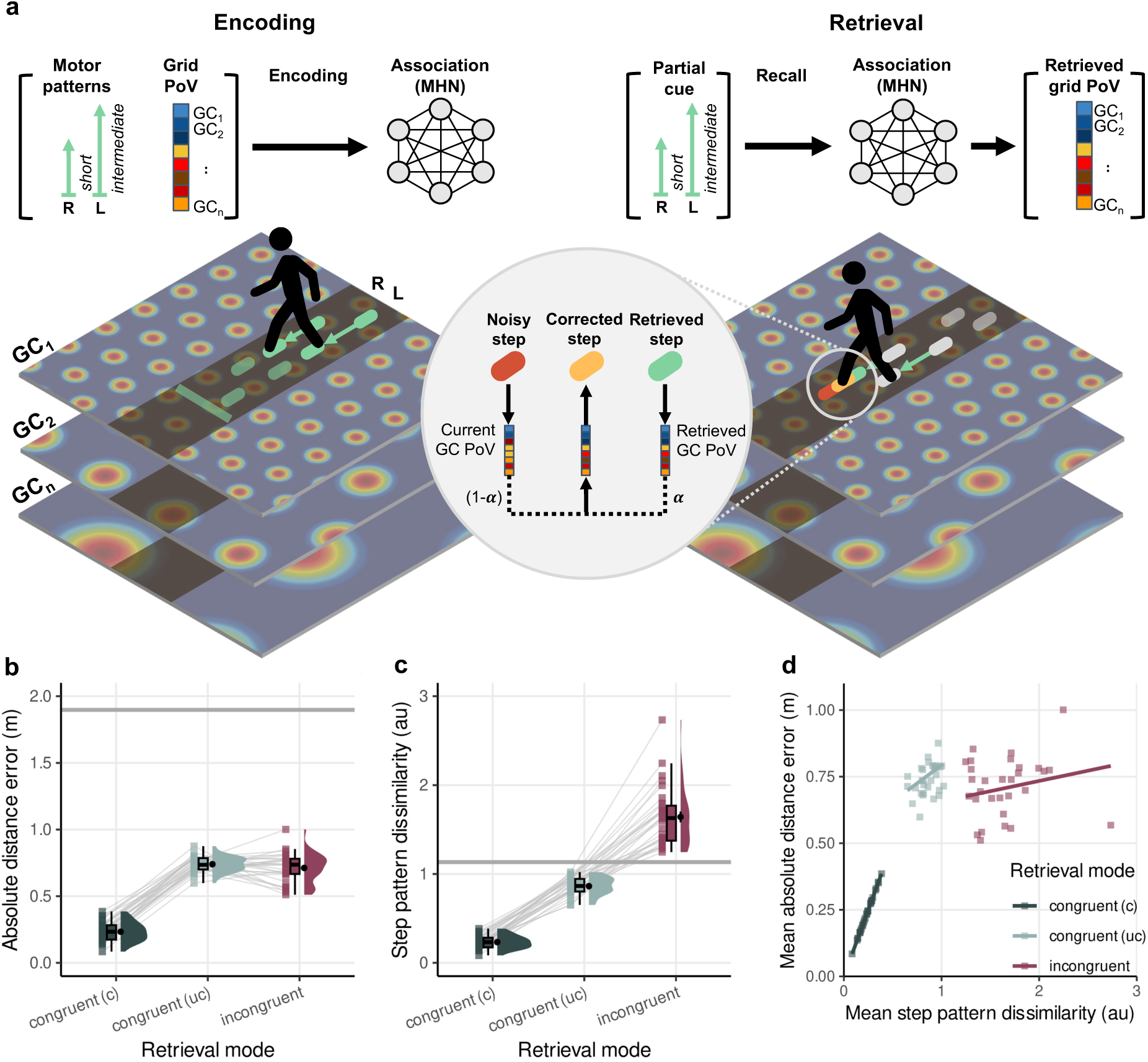
Model schematics and results. **a:** Encoding and retrieval model schematics. During the encoding phase, an artificial agent performs a pseudo-random walk with three distinct target step lengths—small, medium, and long—mirroring the behavioral experiment. At each step along the trajectory, the agent stores a concatenated vector representing motor activity (the current and previous steps, e.g., short and intermediate) along with the corresponding grid cell population vector (PoV) into an autoassociative memory (Modern Hopfield Network, MHN; Ramsauer et al., 2021). In the retrieval phase, vertical and horizontal noise components—defined based on the empirical variance in step lengths—are sampled for each step along the trajectory, corresponding to the random drift of grid cells (Hardcastle et al., 2015). In the congruent retrieval condition, a grid cell population vector is obtained at each position along the noisy trajectory. The agent can retrieve a concatenated population vector of grid cells and motor neurons using a partial cue from the motor neurons (a sequence of two successive steps with the sides of the legs). This retrieved vector contains the encoded grid cell population vector memorized during encoding. A linear combination of the uncorrected (red) and the encoded (green) grid cell population vectors is then calculated; the contribution of each vector can be adjusted by varying the parameter α. The position decoded from this linear combination is considered the error-corrected position of the agent (yellow). **b-d:** Sensorimotor-based error correction in a sample of artificial agents (N = 29; same sample size as in the experiment) mirrors human behavior. **b:** Mean absolute distance error for congruent (with [c] and without error correction [uc]) versus incongruent retrieval. Artificial agents exhibit higher errors during incongruent retrieval compared to congruent retrieval. **c:** Mean step pattern dissimilarity (in arbitrary units) for congruent (with [c] and without error correction [uc]) versus incongruent retrieval. Artificial agents show higher similarity for congruent retrieval than for incongruent retrieval. Violin plots depict the density distribution; boxplots show the median and quartiles; black dots with error bars represent the means ± SEM; colored dots indicate individual data points per condition, and grey horizontal lines refer to the mean chance level (Supplemental Fig. S2). **d:** Relationship between mean step pattern dissimilarity and mean absolute distance error for congruent (with [c] and without error correction [uc]) versus incongruent retrieval. Positive correlations are found for both conditions, with notable differences in intercepts (congruent < incongruent). Dots represent individual agents; lines are based on linear Kendall-Theil regressions.

During the retrieval phase, agents’ walking trajectories were modeled with noise reminiscent of error accumulation observed during rodent and human PI (Durgin et al., 2009; Hardcastle et al., 2015). To define noise in the model, we obtained behavioral step length variability in the empirical data. Specifically, we assumed that participants with poor distance estimation performance during the free retrieval phase might not have formed a reliable long-term representation of the encoded trajectories. Therefore, we draw vertical and horizontal step components from distributions fitted to the free retrieval data filtered for absolute distance errors greater than 2 SD above the sample mean (d_error_ > 0.78). Separate distributions were fitted for each step size, distinguishing between the left and right legs. As a result, the sampled (“uncorrected”) trajectory of a simulated agent incorporates noise that reflects a number of potential sensorimotor and memory-related error sources observed in our sample (cf. Qi & Mou, 2023).

To correct this error, the model retrieves the grid cell PoV’s associated with steps in the stored motor sequence along the noisy trajectory (Fig. 4a, top right). This retrieval from the Hopfield network is triggered by a partial cue - the simulated sensorimotor pattern, yielding the previously encoded grid cell PoV. We then calculate a linear combination of the “uncorrected” and retrieved grid cell PoV’s, where the contribution of each vector can be adjusted by varying the parameter α (Fig. 4a, bottom middle). The position decoded from this linear combination is considered the error-corrected position of the agent.

In contrast, during incongruent retrieval, the agent walks along the trajectory with more regular steps drawn from the previously fitted distribution. Separate distributions were fitted for each trajectory size, distinguishing between the left and right legs. In this scenario, error correction cannot occur due to the ambiguity of the stored keys in the autoassociative memory. Note that since only the target location itself can be accessed during incongruent retrieval, error correction might be expected only at the end of a given trajectory (last step).

Critically, our model results (Fig. 4b-d) mirrored the key experimental observations (Fig. 3d-e, g): agents exhibited lower mean absolute errors under the (corrected) congruent condition compared to the (uncorrected) incongruent condition (Fig. 4b). Similarly, their step pattern dissimilarity (SPD) under the congruent condition was lower than that of the incongruent case (Fig. 4c). Finally, the SPD for the model predicted the mean absolute error (Fig. 4d), consistent with the experimental data. These simulations provide a mechanistic account of how the re-enactment of encoding-related motor patterns can facilitate the retrieval of associated spatial information from memory, leading to error correction and improved accuracy in distance reproduction during PI.

### Accentuated online step correction during congruent retrieval

We proposed an error-correcting mechanism for grid cell drift that may explain the congruency effect observed in our empirical data (see above). Specifically, the model predicts that encoding-congruent motor patterns trigger the correction of upcoming noisy spatial representations throughout a given step sequence, leading to overall better distance reproduction. In contrast, during encoding-incongruent walking, the respective motor patterns do not prompt step-wise error-correction, resulting in poorer distance reproduction. Hence, if neural error correction at least partially manifests behaviorally, congruent retrieval should induce more step corrections than incongruent retrieval.

In an exploratory analysis, we identified sequences where two consecutive steps were taken with the same foot (“correction steps”) and compared their frequency between retrieval modes. Example step patterns containing correction steps are shown in Figure 5a. Indeed, more participants showed step corrections during congruent retrieval (*n* = 22; 76%) compared to incongruent retrieval (n = 15; 52%). In total, between 6% (incongruent) and 10% (congruent) of all sequences contained at least one correction step. These steps were performed a similar number of times with each foot (congruent: 42% left foot; incongruent: 56% left foot) and, on average, appeared to be medium sized (congruent: M = 0.3 m, SD = 0.25 m; incongruent: M = 0.35 m, SD = 0.20 m). A bootstrapped paired *t*-test on a small subset of participants with complete paired data (*n* = 12) suggested a trend for more step corrections during congruent compared to incongruent retrieval (*t*_11_ = 2.52, *P* = 0.08, *d* = 0.73, 95% CI = 0.38 to 1.68; Fig. 5b). Although this effect did not reach statistical significance, likely due to lack of power, it was moderate to strong, indicating that error-correction was more accentuated during congruent walking. Next, we analyzed *where* the correction of steps occurred throughout sequences (i.e. between start = 0 and end position = 1). Based on our model, we expected error-corrections being roughly uniformly distributed during congruent retrieval and highly concentrated at the end of sequences during incongruent retrieval. Consistently, we found that during congruent retrieval, the relative position of correction steps broadly varied throughout sequences, whereas during incongruent retrieval, they clustered at the end of a sequence, i.e. near the end position (Fig. 5c). As a result, the mean position of correction significantly differed across retrieval modes (bootstrapped paired *t*-test: *t*_11_ = -3.33, *P* < 0.05, *d* = -0.96, 95% CI = -2.41 -0.40), which agrees with our model.

**Figure 5.**
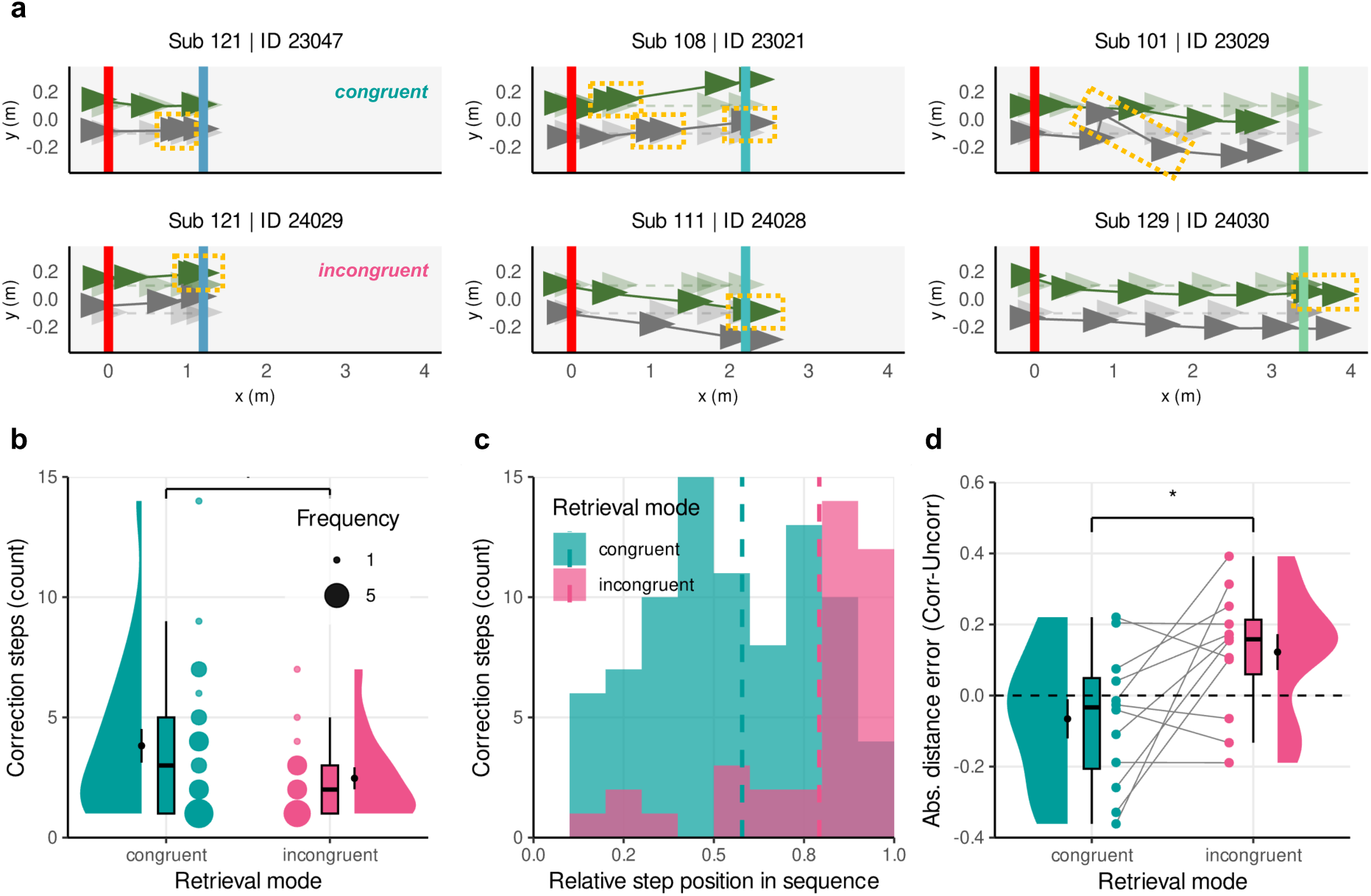
Step correction during congruent and incongruent retrieval. **a:** Step sequences including correction steps (yellow dotted rectangles) of 3 example participants for a short (left), medium (middle) and long (right) target distance trial during congruent (top) and incongruent retrieval (bottom). Participants started from a fixed location (red line at 0 m) and executed left (dark green) and right (gray) steps until they stopped (response). Transparent triangles represent encoded steps towards the target location (blue, cyan, light green lines). **b:** Number of executed correction steps during congruent (green) vs incongruent (magenta) retrieval. Participants tended to perform more correction steps during congruent vs incongruent retrieval (*P* = .08, *d* = 0.73, *N* = 12). Violin plots depict the density distribution, boxplots the median and quartiles, black dots with error bars the means ± SEM, and colored dots individual participants per condition with size of dots representing frequency. **c:** Number of executed correction steps along the linear track (relative position within a sequence) during congruent (green) and incongruent (magenta) retrieval. While during congruent retrieval, correction steps were uniformly executed along the linear track, during incongruent retrieval these steps were more frequent near the response location. Dashed lines reflect the mean relative position within the sequence. **d:** Difference between corrected and uncorrected absolute distance error. Step corrections, on average, reduced error during congruent retrieval (< 0) and enhanced error during incongruent retrieval (> 0). ᐧ *P* < .10, * P < .05.

Ultimately, error-correction is assumed to facilitate distance reproduction. If true, participants’ absolute distance error should be higher without correction steps. To simulate this unobserved scenario, we subtracted the sum of corrections steps per sequence from the recorded end location (response) and calculated the difference between “corrected” and “uncorrected” absolute distance error for each participant. A bootstrapped paired *t*-test on difference values indicated a significant effect of retrieval modes (*t*_11_ = -2.58, *P* = 0.04, *d* = - 0.75, 95% CI = -1.43 to -0.30; Fig. 5d), suggesting a distinct influence of correction steps on distance reproduction. On average, correction steps non-significantly improved distance reproduction by 4.3 cm during congruent retrieval (bootstrapped paired *t*-test: *t*_21_ = -0.81, *P* = 0.44, *d* = -0.17, 95% CI = -0.74 to 0.22) and significantly impaired distance reproduction by 12 cm during incongruent retrieval (*t*_14_ = 3.20, *P* < 0.01, *d* = 0.83, 95% CI = 0.38 to 1.59). As expected, uncorrected distance reproduction was comparable between congruent and incongruent retrieval (*t*_11_ = -0.40, *P* = 0.69, *d* = -0.12, 95% CI = -0.89 to 0.46). Together, these exploratory analyses imply some interesting similarities between the model-predicted error corrections in grid cells and the step corrections observed in human behavior.

## Discussion

This study addressed the role of motor re-enactment in preserving an accurate distance estimate during path integration (PI) despite error accumulation (Hardcastle et al., 2015) and without vision. Building on the role of sensorimotor processes in human memory (e.g. Ianí, 2019), we show that participants who spontaneously or intentionally re-enacted the irregular step patterns learned at encoding reproduced distance more accurately, whereas suppressing re-enactment with a regular gait impaired performance. Critically, we demonstrate that the benefit of matching movement patterns aligns with a neurocomputational model in which retrieval-related noise in grid cell activity is corrected through the reactivation of encoding-related motor activity. This suggests that factors other than environmental input can help correct for drift in the PI system. Finally, our observation of corrective steps in the behavioral data hints at a potential behavioral manifestation of this neural correction mechanism.

By leveraging detailed movement analysis, we identified pronounced similarities between the step patterns participants used during encoding and their freely chosen gait during retrieval. This observation not only confirms the involvement of spatial and motor sequence memory in the re-enactment of the number, length and order of steps, but also reveals a training-induced preference for performing complex spatio-temporal step patterns over more energy-efficient gait patterns, typically observed during real-world locomotion (e.g. Donelan, Kram & Kuo, 2001; Seethapathi & Srinivasan, 2015; Selinger et al., 2019). Given the limited number of trial repetitions (16 per target) up to this point, this re-enactment is unlikely attributable to habitual motor learning, as even simple motor skills require excessive rehearsal to become automatic (e.g. Immink, Verwey & Wright, 2020; Kami et al., 1995; Moors & de Houwer, 2006; Puttemans, Wenderoth & Swinnen, 2005). Instead, post-experimental debriefings indicated that participants consciously attempted to mimic the learned step patterns, implying a functional role in non-visual distance estimation.

Our data revealed that matching step patterns across encoding and retrieval results in better distance reproduction than adopting a regular, encoding-incongruent gait. Critically, participants performed above chance even during incongruent retrieval where reliance on motor sequence memory alone was not feasible, suggesting that they encoded integrated spatial information in addition to sequence information. Given the task’s strong reliance on idiothetic cues, this finding suggests that locomotion-induced variations in body signals may alter the perception of distance. Indeed, studies have shown that changing the type of repetitive gait across outbound and return paths can degrade distance reproduction (Abdolvahab et al., 2015; Harrison, 2020; Harrison & Davis, 2023; Harrison et al., 2021; Mittelstaedt & Mittelstaedt, 2001; Schwartz, 1999; Turvey et al., 2009, 2012). While attempts to characterize the biological odometer by simple locomotion parameters like the number or length of steps, elapsed time or energy consumption remained inconclusive (Durgin et al., 2009; Mittelstaedt & Mittelstaedt, 2001; Schwartz, 1999; Turvey et al., 2009; Wittlinger, Wehner, & Wolf, 2006, 2007), its reliance on higher-order idiothetic classes such as gait symmetry (Abdolvahab et al., 2015; Turvey et al., 2009, 2012), action mode (Chrastil & Warren, 2014) or reference frame (Harrison, 2020; Harrison & Davis, 2023) has been proposed. For instance, Turvey et al. (2009) reported indistinguishable accuracy for forward-forward and backward-forward walking, possibly because the same symmetry class is shared by both gaits. In contrast, our data revealed a more gradual relationship: even subtle deviations from encoded movements incrementally reduced reproduction accuracy, suggesting the degree of idiothetic input similarity predicts performance.

The reliance on movement similarity for accurate distance estimation further aligns with well-established principles of memory emphasizing the importance of encoding-retrieval overlap. Frameworks like *transfer-appropriate processing* (Morris, Bransford, & Franks, 1977), *context-, state-, and mood-dependent memory* (Godden & Baddeley, 1975; Eich, 1980; Eich & Metcalfe, 1989), and the *encoding specificity principle* (Tulving & Thomson, 1973), posit that memory retrieval is optimized when encoding and retrieval conditions match. In our task, the “condition” reflected a unique motor pattern that was directly tied to spatial information, forming an associative memory trace. By re-enacting the original motor pattern, participants maximized the conditional overlap between encoding and retrieval, reinforcing the reinstatement of spatial information. This interpretation is consistent with converging evidence suggesting that memories are grounded in sensorimotor activity patterns reactivated during retrieval (Danker & Anderson, 2010; Ianí, 2019; Kent & Lamberts, 2008; Masumoto et al., 2006; Meyer et al., 2024; Nilsson et al., 2000; Nyberg et al., 2001) and re-enacting bodily states enhances memory (Dijkstra et al., 2007; Iverson & Goldin-Meadow, 1998; Morse et al., 2015; Morsella & Krauss, 2004; Ping et al., 2014; Wynn et al., 2019). By demonstrating the importance of full-body movements for spatial memory representations, our data, together with reports of hippocampal involvement in motor memories (Albouy et al., 2015; Döhring et al., 2017; Jacobacci et al., 2022), challenges the classic distinction between declarative and procedural memory (Eichenbaum & Cohen, 2004; Gabrieli, 1998; Squire, 1992) and underscores the embodied nature of memory (Barsalou, 2008; Dijkstra & Post, 2015; Ianì, 2019).

To mechanistically explain our behavioral findings, we developed a neural-network model that links motor memory to grid cell coding. Building on diverse evidence showing that pure self-motion integration is inherently unstable (Burak & Fiete, 2009; Hardcastle et al., 2015; Hasselmo, 2008; Hasselmo & Brandon, 2012), our model assumes that noise displaces the grid code. It then proposes that encoding-congruent motor activity can serve as a corrective signal to mitigate noise-induced drift within the grid cell population. Whereas earlier work relied on environmental cues to reset accumulated error (Evans et al., 2016; Hardcastle et al., 2015; Pollock et al., 2018), our simulations highlight the corrective power of body-derived cues in complete darkness. To our knowledge this is the first model showing that motor information alone can stabilize PI estimates, revealing a potentially overlooked contribution of the motor system to spatial accuracy, particularly without stable environmental cues. Anatomical data across species support the notion that motor information reaches the EC (Burwell & Amaral, 1998; Insausti, Amaral & Cowan, 1987; Sepulcre et al., 2012; Reznik et al., 2024).

The model successfully simulates the observed behavioral pattern, demonstrating reduced distance error in agents employing motor-driven error correction compared to those without. Furthermore, we observed a trend for participants to exhibit more corrective steps during congruent retrieval than during incongruent retrieval, potentially reflecting a behavioral manifestation of the proposed correction mechanism. Although this observation aligns with our model’s predictions, no statistically significant benefit of these step corrections was found for distance reproduction. One plausible reason for the lack of significance is low power resulting from analyzing only a small subsample of participants together with the coarse definition of correction steps that includes very large steps resulting from postural imbalance. Another possibility is that online error correction operates primarily at a neural level, not always requiring overt behavioral adjustments. Future research with appropriate statistical power and more sensitive behavioral measures could directly address the role of corrective motor behavior in path integration. Moreover, employing modern mobile neuroimaging techniques, such as optically pumped magnetometers (Boto et al., 2018) or intracranial recordings (Stangl, Maoz & Suthana, 2023), will be crucial for directly investigating the role of sensorimotor reactivation and motor-driven correction of path integration errors.

In conclusion, we provide evidence for a functional link between consistent irregular movement patterns and accurate distance estimation during self-motion. We demonstrate that re-enacting these patterns supports distance reproduction, while preventing re-enactment impairs performance. Our neurocomputational model provides a mechanistic account for this phenomenon, suggesting that encoding-congruent motor activity can mitigate noise in grid cell representations during retrieval. This constitutes the first model of path integration correction via motor representations. These findings thus highlight the crucial role of embodied sensorimotor processes in spatial cognition, extending previous research on haptic distance perception and aligning with established theories of memory retrieval.

## Methods

### Participants

34 healthy younger adults took part in the study, all recruited from the internal database of the Max Planck Institute for Human Cognitive and Brain Sciences, Leipzig, Germany. The sample size was estimated *a priori* by a power analysis using G*Power version 3.1.9.6 (Faul, Erdfelder, Buchner & Lang, 2009). This yields a sample size of 28 participants to ensure a statistical power of 95% (*f^2^* = 0.25, α = 0.05, repeated measures ANOVA, two-sided test). To account for drop-out, we recruited 6 additional participants. *Post hoc*, 3 participants were excluded due to poor task performance that was classified if the variability in signed distance reproduction errors was greater than 2× the interquartile range compared to the upper quartile of sample mean errors. Another participant was excluded due to data loss resulting from technical difficulties during the experiment. Accordingly, our final sample comprised 29 participants (12 female, 14 male, 3 non-binary; age: M = 26.86 years; SD = 4.45 years, range = 19-35 years). During the time of data acquisition, all participants reported normal or corrected-to-normal vision, no current or history of neurological, psychiatric or motor disorder and being right- handed. Supplementary Table S1 summarizes group-specific sample characteristics. In order to limit the individual variability of step sizes, we recruited only participants with a body height ranging 164-186 cm (Supplementary Tab. S2 for a full record of body measures and demographics). All participants gave written informed consent after being instructed about the general goals and procedures of the study. All proceedings were approved by the regional ethics committee of the Medical Faculty at the University of Leipzig following the ethical recommendations laid down in the Declaration of Helsinki. Participants received an expense allowance of €12 per hour.

### Apparatus

We combined state-of-the-art immersive virtual reality (VR) and motion capture (MoCap) technology (Supplementary Fig. S1a) to control step execution through dynamic visual stimulation while allowing for naturalistic full-body movements. Participants were immersed in virtual environments built with Unity3D/C# (version 2021.3.3f1) and presented on a Pico Neo 3 Pro Eye Head-Mounted Display (HMD; 3664 × 1920 pixels, field of view: 90-98 degrees, refresh rate: 90 Hertz, weight: 620 grams; Pico Interactive, San Francisco, CA, US; https://www.picoxr.com). For sub-millimeter precise tracking of body kinematics, we used a Vicon MoCap system (Vicon, Oxford, UK; https://www.vicon.com/) incorporating 9 Vero v2.2 high-resolution infrared cameras (2.2 megapixel at 330 frames per second) mounted on a ceiling frame to cover a tracking volume of approximately 4 × 5 × 2.5 meters. Within this space, three-dimensional motion data (sampled at 200 Hertz) was captured from passive retro- reflective markers attached to 14 rigid body patches (Supplementary Fig. S1b), the HMD, and a handheld motion controller. The motion data was live-streamed via the Vicon system’s PoE switch using Vicon Tracker software, hosted on a stationary computer (AMD Ryzen 5 3600 6- core 3.6-4.2 gigahertz processor, NVIDIA GeForce RTX 3090 graphics card, Intel I350 network card, running Microsoft Windows 10). Simultaneously, VR images (game frames) were broadcasted to the head-mounted display (HMD) display via the Pico Link client using a Wi-Fi 6 router (ASUS RT-AX82U). This setup ensured a wireless VR experience with separate data streams for motion tracking and VR rendering. Debriefing forms, completed after the experiment, were implemented as one custom-made Unity3D application presented via standard computer monitor (2560 × 1440 pixels).

### Virtual environments

A one-dimensional linear track (width: 0.5 m, length: infinite) projected onto a boundless ground plane within a light sky box constitutes the virtual main environment (Fig. 1c). Both the track and the ground were given a texture to aid optical flow during locomotion, with fine-grained tiles that could not be used as local cues. No local or distal landmarks were provided, forcing participants to focus on self-motion cues in order to keep track of their own position relative to a fixed starting position. To enhance the subjective experience of immersion and virtual body ownership (Waltemate et al., 2018), we animated a humanoid virtual avatar (Supplementary Fig. S1c) and mapped it onto the body of each participant. Body parts of the avatar were scaled individually based on body measurements taken prior to the experiment (Supplementary Tab. 2). Task interactions were performed via the controller, carried in the right hand. Instructions and feedback were presented on a display attached to the controller position (Supplementary Fig. S1d).

At the beginning of the experiment and in-between trials, participants were presented with a virtual base environment consisting of an infinite dark ground plane embedded within a light sky box, again without any local or distal landmarks.

### Distance reproduction task

We developed a novel immersive VR-based distance reproduction task that required participants to learn the location of three targets, located at different distances (short = 1.2 m, medium = 2.2 m, long = 3.4 m) from an invariant starting location. Since the environment contained no distal or local cues, participants had to rely on the traveled distance based on self-motion cues to succeed in the task. The task consisted of a training and test phase. Videos of example trials for each phase can be found on the Open Science Framework (https://osf.io/dhe25/).

During the training phase, encoding and immediate retrieval trials were alternated. Participants iteratively learned the locations of the targets along the linear track, by adopting unique leg movements. In each encoding trial, participants were asked to follow a predefined step pattern by moving their feet onto successively appearing footholds (flat, monochromatic rectangles of approximately 34 × 24 centimeters) along the linear track. Step patterns were made of three step lengths (small = 0.2 m, intermediate = 0.4 m, large = 0.6 m) concatenated in random order, so that the number of steps per length was fixed for each target distance. For each step pattern, the time intervals in between steps (range: 0.1-2 s) were randomly sampled across steps and different for each participant. Right and left footholds appeared alternately with 46% of patterns starting with the left foot and disappeared after a step was detected. Some characteristics of the pre-defined step patterns are summarized in Supplementary Table S4. At the end of each step pattern, the location of the target was indicated by a uniquely color-coded line, i.e. a target’s identity, marked on the ground along the linear track. The assignment of identities (red, green, blue) to target distances was randomized across participants. On arrival, participants were instructed to memorize the target location and confirm by pressing a button before the beginning of the next trial. In the subsequent immediate retrieval trial, the participants were instructed to navigate to a given target location by walking in a straight line in complete darkness. As vision was unavailable, the reproduction of path length purely relied on body-based (i.e. idiothetic) self-motion cues including vestibular and proprioceptive feedback as well as motor efference copies. Importantly, we gave no instruction on how to walk, thus allowing for intrinsic coordination of body movements. On arrival, participants were asked to confirm their distance estimate by pressing a button before being teleported into the virtual base environment where they received coarse visual feedback on their accuracy in the form of one of five smiley faces. These smileys were based on the linear displacement of responses relative to target locations (≥ 1 m = very bad, < 1 m = bad, < 0.5 m = intermediate, < 0.25 m = good, < 0.125 m = very good). No information about the error’s sign (over-/undershoot) was provided and the correct target was not displayed again.

During the test phase, participants had to navigate back to the targets with the HMD display was blanked under three conditions (‘free retrieval’, ‘congruent retrieval’, ‘incongruent retrieval’), which varied in terms of instructions related to the type of locomotion (for trial order and block description see “Methods” section under “Experimental protocol”). Free retrieval trials were similar to immediate retrieval trials during training, but without receiving feedback at any time. No instructions were given on how they should move, thus distance estimates in these trials served as the central memory read-out resulting from previous learning. In case of enquiry, we specified that one should walk in a self-paced way, so that the required distance is optimally reproduced. Here we hypothesized that if the participants’ encoding-related body movements play an effective role in their path estimations, they should tend to intuitively re- enact them, even if they have the choice to use a different (e.g. regular and thus more energy-efficient) step pattern. In contrast, in congruent and incongruent trials, we explicitly specified how participants should move to reach the target. In congruent trials, participants were instructed to precisely mimic the step pattern previously followed during the training phase. In incongruent trials, they were asked to use self-paced regular steps of the same length in order to reach the target. These two conditions allowed us to examine how a (mis)match in sequential body movements between encoding and retrieval affects distance reproduction. After each trial of the test phase, we additionally asked participants to rate on how certain they were about each response by adjusting a slider (originally positioned at the center of the slider) on a subjective scale ranging from “very certain” to “very uncertain” printed on the text display attached to the handheld controller (Supplementary Fig. S1d).

Before the beginning of each trial of the training or test phase, participants were teleported into the virtual base environment and asked to collect a virtual ball (radius = 2.5 cm) floating above the ground at a pseudo-random location. By navigating to the ball, they performed a short random walk aiming to reset their path integration estimate of the previous trial. Once the ball was collected via button press, a red line and arrow appeared on the ground, marking the start location and orientation of the upcoming trial. Participants aligned their feet relative to the line and their head in an upright position by fixating on a distant virtual cross. The subsequent transition from the base to the main virtual environment was triggered by button press.

### Procedure

A single experimental session was divided into three parts - preparation, main task and debriefing - and lasted 2-3.25 hours (M = 2.68, SD = 0.32) in total, of which participants spent 1.25-1.75 hours in VR (M = 1.51, SD = 0.14).

### Preparation

After giving participation consent, the experimenter took several body measures (body height, arm span, inseam length, hip width and arm length), attached all rigid body marker patches to the participants body, and carried out the individualized avatar mapping. For the whole experiment, participants did not wear shoes.

During a familiarization period, participants practiced a minimum of two trials of encoding and immediate retrieval with target identities, distances and step patterns that were different from those used during the main task. These trials aimed to habituate the participants with the virtual environments and to minimize potential ambiguities regarding the task instructions. To avoid potential confusion, the linear track and ground plane during familiarization were of a different color to those used during the main task, signaling different contexts.

### Main task

Next, participants completed the distance reproduction task (see “Methods” section under “Distance reproduction task”), for which we used a 3 × 3 within-subject design (Fig. 1a-b) with two factors: target distance (small, medium, long) and retrieval mode (free, congruent, incongruent). During the training phase, there were 10 iterations of encoding and immediate retrieval trials, organized in randomized mini-blocks of three trials. For each mini-block we sampled each target distance once in a way that no consecutive trials sampled the same target distance. During the test phase, participants completed three blocks with six repetitions per target distance in random order. Each block corresponded to one of three conditions (free retrieval, congruent, incongruent). The test phase always started with free retrieval trials, whereas the order for congruent and incongruent retrieval blocks was randomized across participants (congruent first: 52% of participants). Walking directions of successive trials were rotated by 180 degrees and randomized across target distances to prevent the use of location-related cues other than body-based cues (e.g. ambient noise).

### Debriefing

After completing the main task, participants underwent a series of short tasks and questionnaires on a standard computer screen (see “Methods” under ”Apparatus”). First, we presented participants with a bird’s eye view of the linear track spanning the full-width of the screen (left-to-right side) with the start location being displayed as a vertical line. Along this track, we asked them to consecutively place separate markers of target distances. To do so, they needed to drag-and-drop vertical tags (i.e. the target ID) onto the track in order to indicate their respective abstract place memories. In addition, we asked participants about explicit encoding-related knowledge such as the number steps and target distances. Second, we asked various questions about task-related strategies, body attention, volition, postural control, motion sickness, motivation. Descriptive results of the debriefing are summarized in Supplementary Table S4.

### Data processing and analysis

#### Distance reproduction performance

The accuracy of estimating and reproducing distances from memory was determined by assessing the deviation from the true target distance. For each trial, we subtracted the target distance between the start and the target location from the reproduced distance between the start location and the center of the right and left toes at the time of the response (button press). While the *absolute distance error* captures the unsigned magnitude of deviations, the *signed distance error* indicates the direction of deviation, i.e. overshooting and undershooting, with its variability over trials designating precision (Fig. 1d-f).

To assess training performance in immediate retrieval trials (Fig. 1h-i), we tested the effect of repetition and target distances on mean absolute distance errors using a bootstrapped 2-way repeated measures ANOVA with *post hoc* bootstrapped paired *t*-tests. Deviations from 0 in mean signed distance errors were tested using bootstrapped 1-sample *t*- tests.

Successful learning was verified by comparing the mean absolute distance errors in free retrieval trials to target-specific mean chance levels using bootstrapped 1-sample *t*-tests (Fig. 1j-k). The chance distributions were generated by uniformly sampling random responses along the linear track, spanning 0.05 to 6.5 meters from the start (Supplementary Fig. S2a, left), with the maximum response distance of 0.5 meters from a physical boundary (table or wall). Simulated distance errors were calculated for 1,000 artificial participants, each following the original trial structure of 6 trials per target distance, yielding a total of 18,000 random responses. The resulting chance distributions of distance errors for each target distance are reported in Supplementary Fig. S2a, left. Furthermore, we investigated the variability in distance reproduction over trials with respect to the target distance (Supplementary Fig. S3b). For each participant we computed the coefficient of variation (CV) as the standard deviation σ̄ of the log-transformed distances *d* by applying the formula (Durgin et al., 2009): *CV* = *exp*(σ̄(*log*(*d*)) − 1). To verify Weber’s law, differences in the CV between target distances were tested using a bootstrapped 1-way repeated measures ANOVA with *post hoc* bootstrapped paired *t*-tests.

#### Movement characteristics

##### Preprocessing

Raw motion data was recorded for each rigid body (i.e., marker plates, HMD and motion controller) at a rate of 200 Hz. Signal dropouts within sequences were corrected using linear and spherical interpolation. To estimate precise joint positions and angles, we applied the inverse kinematic solver *UltimateIK* (https://github.com/sebastianstarke/AI4Animation) to fit a Skinned Multi-Person Linear Model of the human body (Loper et al., 2015) to each participant’s body measurements. The 3D model was animated frame by frame based on the recorded rigid body motions. Subsequently, the data were downsampled to 100 Hz, and motion sequences were filtered to include only the period between the confirmation of the start position (button press on the controller) and the confirmation of the target position (button press at the end of an encoding trial) or its reproduction (button press at the end of a retrieval trial).

##### Step detection

A single step was considered to start when the velocity of a given foot exceeds the 200 mm/s threshold. The step ends as soon as it falls below the threshold again. To avoid false detections, a Butterworth low-pass filter was applied to the foot velocities and detected steps with a positive stride length (i.e., the distance between two ground contacts of the same foot) shorter than 0.05 m and longer than 1 m were excluded for the analysis of non-corrective steps.

##### Number of steps

We counted the number of executed steps during both encoding and retrieval and calculated the difference between phases (Fig. 2b). Bootstrapped 1-sample *t*-tests were performed to test the deviation from a perfect match (i.e. difference of 0), separately for each target distance and the average. Differences across target distances were assessed using a bootstrapped 1-way repeated measures ANOVA.

##### Length of steps

The step length was determined at the end of a completed step by measuring the Euclidean distance between the grounded opposite toes (support phase) projected onto the 2D walking direction, i.e., the linear vector from the start position to the participants’ final position at the end of a trial. In cases where two or more steps were taken with the same foot in succession (“correction steps”), we only considered the step lengths of corrected forward movements, so that step lengths within a sequence add up to the final end position (i.e. the distance estimate).

##### Number of step length clusters

During encoding, participants encoded step patterns consisting of 3 step lengths. To investigate whether this number was evident also in freely retrieved step patterns, we performed a cluster analysis for each participant, considering both all step lengths (pooled) and grouped by target distance (Fig. 2c-d). First, we estimated the density function of step length using Kernel Density Estimation (KDE) with the bandwidth individually selected through unbiased cross-validation. Local maxima in the KDE, representing potential clusters, were identified as peaks exceeding 20% of the maximum density value (Fig. 2c). To ensure that each peak was distinct, any adjacent peaks within 0.05 m of one another were consolidated, retaining only the most prominent peak in such cases. The number of significant KDE peaks per participant was quantified as the number of clusters (Fig. 2d, top). We aggregated cluster counts across participants and calculated the difference to the ground truth of 3 encoded step lengths (Fig. 2d, bottom). Deviations from the ground truth were tested using bootstrapped 1-sample t-tests against the null hypothesis of 0 difference. A bootstrapped 1-way repeated measures ANOVA was used to evaluate variations across target distances.

##### Step length class assignment

Building on the mean cluster count from the preceding analysis, step lengths were classified into three distinct categories: small (0.2 m), intermediate (0.4 m), and large (0.6 m). For each participant, we assigned both pooled (Fig. 2e, top) and target-specific step lengths to these classes using Jenks Natural Breaks Classification (Jenks, 1967), a method that optimally partitions 1D data by minimizing within-class variance and maximizing between-class variance. Participants with fewer than two unique step lengths were excluded to ensure reliable class assignments. For each step per target distance, the deviation from the designated target cluster step length was calculated, providing a measure of accuracy for each cluster (Fig. 2e, bottom). To evaluate whether these deviations significantly differed from 0, we used bootstrapped 1-sample *t*-tests. Additionally, a bootstrapped 2-way repeated measures ANOVA was conducted to investigate the effects of step length class and target distance.

##### Step pattern dissimilarity

The step pattern dissimilarity (SPD) quantifies the discrepancy between the encoding-related step pattern (“desired”) and the retrieved steps (“performed”; Fig. 1g). Since executed step patterns may vary in the number of steps and their lengths, we used Dynamic Time Warping (DTW) to match the executed steps separately for each side (left and right foot) with the steps in the target pattern, ensuring that no step is matched twice. The DTW algorithm (Giorgino, 2009) aligns two step sequences by non-linearly stretching or compressing them to minimize their temporal differences, enabling comparison, even if they are out of phase or differ in length. The SPD was formalized as:

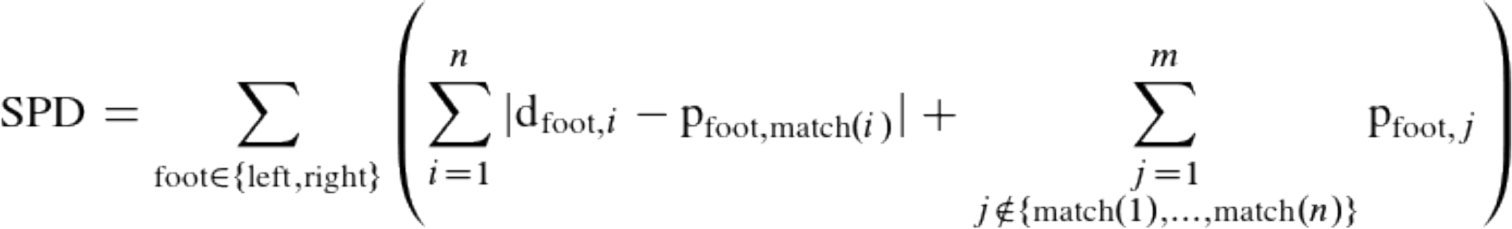

where *d*_foot,_*_i_* is the *i*-th desired step length, *p*_foot,_*_j_* is the *j*-th performed step length, and the function match(*i*) identifies the index of the most similar performed step length to the desired step at index *i*, based on the DTW algorithm. The first summation within brackets computes the absolute step length difference for each matched step pair among *n* desired steps. The second summation adds the full lengths of any extra or missed steps among *m* performed steps. Finally, the values determined for the left and right sides are summed. A SPD value equal to 0 reflects a perfect replica of encoding-related steps in terms of their number, length, and order. Similar to absolute distance errors (see above), we determined a simulated chance level for SPD values by focusing on arbitrarily ordered steps. For each target distance and the average, we generated random step patterns uniformly sampled between the start and target locations (Supplementary Fig. S2b, left), ensuring the same number of steps as during encoding. Since participants’ step patterns could vary in regularity (depending on how well the step patterns were learned and re-enacted), we introduced a step sample width as a regularity index ranging from 0 to 1, with 0 reflecting the target distance divided by number of steps (maximum regularity). For example, with a step sample width of 0.5, steps were uniformly sampled within a 50 cm range around the regular step length (see Supplementary Fig. S2c). The step sample width was divided into 10 bins and evenly distributed across 1,000 artificial participants, each retaining the empirical trial structure. This approach ensured that the chance distribution accounted for varying levels of step regularity. To evaluate whether empirical SPD values fell significantly below chance (Fig. 2f), we performed bootstrapped 1- sample *t*-tests against the mean chance level for each target distance and the average. The SPD chance distributions are shown in Supplementary Figure S2b, right. Differences across target distances were assessed using a bootstrapped 1-way repeated measures ANOVA with *post hoc* bootstrapped paired *t*-tests. The predictive power of SPD values in explaining variance in absolute distance errors (see above) was evaluated using Kendall-Theil regressions, separately performed for each target distance as well as for the averaged data (Fig. 2g).

### Congruent versus incongruent retrieval

#### Movement characteristics

To verify that participants followed the task instructions (i.e. either mirroring the encoding step pattern or using a self-paced regular step length), we compared both the within-sequence variance of step lengths (Fig. 3b) and SPD values (Fig. 3d) between congruent and incongruent retrieval using a bootstrapped paired *t*-test. In addition, SPD values were tested against the average chance level (as described above) using bootstrapped 1- sample *t*-tests, one for each retrieval mode. On a more fine-grained level, we assigned each produced step length to one out of three target step lengths using DTW and subsequently estimated the effect of target step length and retrieval mode via a bootstrapped 2-way repeated measures ANOVA (Fig. 3c). For one participant, all produced steps classified as small (0.2 m) were smaller than 0.05 m and therefore excluded.

#### Distance reproduction

Differences in the mean absolute distance error between retrieval modes were evaluated using a bootstrapped paired *t*-test (Fig. 3e). To assess the deviation from the average chance level, individual bootstrapped 1-sample *t*-tests were conducted. Lastly, a Kendall-Theil regression tested the predictive effect of SPD values on absolute distance errors for each retrieval mode (Fig. 3g).

#### Corrections steps

We filtered sequences in which the same foot was used for two consecutive steps, referred as “correction steps”. For each retrieval mode, we descriptively report 1) how many participants exhibited at least one correction step, 2) the number of sequences containing correction steps relative to all sequences, and 3) the proportion of corrections steps executed with the left vs right foot. To test differences in the total number of performed correction steps, we employed a bootstrapped paired *t*-test (Fig. 5b). We then calculated each step’s position along the linear track relative to the maximum step position within a sequence and determined the mean relative position for each retrieval mode (Fig. 5c). To assess differences between congruent and incongruent retrieval, we conducted a bootstrapped paired *t*-test. Finally, we explored the impact of step correction on distance reproduction performance (Fig. 5d). To do this, we computed “uncorrected” distance estimates by subtracting the sum of correction step lengths within a sequence from the reproduced distances per trial and calculated the corresponding absolute distance error (as described above). We subsequently quantified the difference in absolute distance error between “corrected” and “uncorrected” estimates and compared this difference between congruent and incongruent retrieval modes using a bootstrapped paired *t*-test. Deviations from 0 (no difference) were tested using individual bootstrapped 1-sample *t*-tests. Note that the number of participants displaying correction steps differed between congruent and incongruent retrieval modes. Therefore, statistical comparisons across retrieval modes were performed only on a subsample with complete paired data.

### Statistical analysis

All statistical analyses and visualizations were conducted in R 4.4.0 using RStudio developer environment (version 2023.06.0). We employed custom bootstrap-based functions to perform repeated measures ANOVA and *t*-tests, thereby enhancing the robustness and reliability of our statistical inferences by mitigating the assumptions inherent in traditional parametric tests. To generate null distributions, we resampled residuals excluding the primary effects of interest for ANOVAs, resampled the differences between centered observations for paired *t*- tests, and resampled the entire centered dataset for 1-sample *t*-tests. All resampling distributions were based on 10,000 iterations. These bootstrap samples facilitated the construction of distributions of test statistics, which were then used to derive *P*-values by comparing empirical values against their bootstrap counterparts. Additionally, we calculated bootstrapped 95% confidence intervals for effect sizes (Cohen’s *d* for *t*-tests and partial eta squared for ANOVAs) using the percentile method. To assess the linear relationship between two variables, we utilized Kendall-Theil estimators provided by the *mblm* package in R (https://CRAN.R-project.org/package=mblm), a robust and assumption-free approach for fitting regression lines. Efron’s R-squared was reported as the measure of effect size. All analyses were conducted using two-sided tests with an alpha level of 5%. To control for the risk of Type I errors due to multiple comparisons, we applied the Bonferroni correction.

#### Modeling

##### Grid Cells

We modelled grid cells as firing rate maps, where each (x, y) location corresponds to a physical position in space and provides the firing rate of all neurons (Bicanski and Burgess, 2018). Grid cell maps were organized into modules that differ in frequency, scaling offsets, and orientations. The maps are matrices of 600 × 600 pixels, created by calculating cosine waves with a phase offset of 60 degrees as described below.

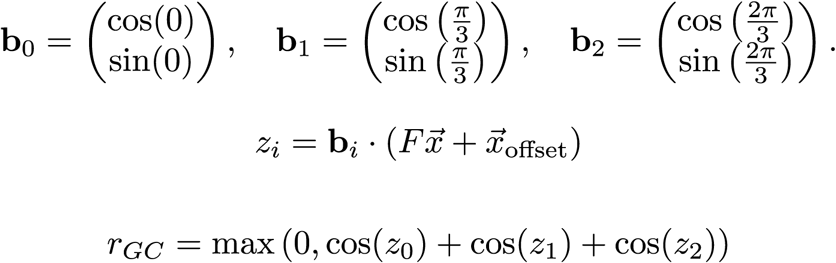

Here, b_0_, b_1_, b_2_ are the unit vectors of the cosine waves, F is the module frequency, and x_offset_ is sampled from values along the principal axes formed by two adjacent equilateral triangles within the grid. By selecting x_offset_ along these axes, we uniformly covered the entire space with grid cell spatial fields (Hafting et al., 2005) – i.e. each rhomboid that was bordered by 4 grid fields was covered with equally spaced offsets. Six values were sampled uniformly along one axis, resulting in 36 different x_offset_ values. We used seven modules with 36 cells in each.

#### Modern Hopfield Network

A classical autoassociative memory model is the Hopfield network (Hopfield, 1982). We used a Modern Hopfield network with continuous-value storage (Krotov & Hopfield, 2016, 2020; Ramsauer et al., 2020). This version stores approximately 2^d/2^ patterns instead of about 0.14d (Demircigil et al., 2017). Patterns are retrieved from the stored matrix by iteratively applying the following update rule until convergence:

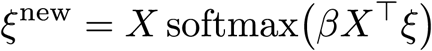

In this equation, ξ^new^ is the retrieved state pattern, X is the matrix of stored patterns, β is the inverse temperature parameter, and ξ is the pattern from the previous iteration. After several iterations, the pattern converges to a stable attractor state. Higher inverse temperature reduces the probability of metastable states, where the system is partially settled between multiple states. The weights are stored in the matrix X, using a learning process equivalent to Hebbian learning.

We chose the Modern Hopfield Network due to its ability to store exponentially many memories with low interference between states.

#### Error-correction

During encoding, we generated pseudo-random trajectories composed of steps with three lengths (0.2 m, 0.4 m, and 0.6 m) for three target distances (1.2 m, 2.2 m, and 3.4 m), similar to the experimental conditions. At each step, we formed a tuple from the current and previous steps and their corresponding leg sides (e.g., [leg_side_t_, step_t_, leg_side_(t-1)_, step_(t-1)_]) and mapped it to a binary vector representing a motor pattern.

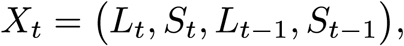

We mapped tuples to binary vectors so that they remain orthogonal in the mapped space.

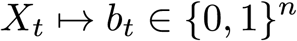

We concatenated this motor pattern with a grid cell population vector obtained from the grid cell frequency maps.

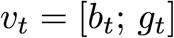

We then encoded this combined vector into the Modern Hopfield Network.

For retrieval, we can differentiate between congruent and incongruent cases. In the congruent case, we fitted the distributions of the x and y step components to data from the free retrieval phase for trajectories with high error (d_error_ > M + 2SD; M = 0.28, SD = 0.25). We estimated separate distributions for each leg side and step size combination (left 0.2 m, right 0.2 m, left 0.4 m, right 0.4 m, left 0.6 m, right 0.6 m). Our hypothesis was that these participants performed poorly in distance estimation because they failed to form a reliable long-term representation of the trajectories. The sampled uncorrected steps should accumulate error along the trajectory, analogous to the random drift of grid cells (Hardcastle et al., 2015). The motor pattern b_t_ is used for retrieval of the concatenated vector v_t_ containing both the encoded motor and grid cell vectors (b_t_ and g_t_). Thus, we had access to both the current and the encoded grid cell population vectors.

We applied a linear correction to the current grid cell population vector g_curr_ using the stored encoded vector g_enc_. This correction is modulated by *α*:

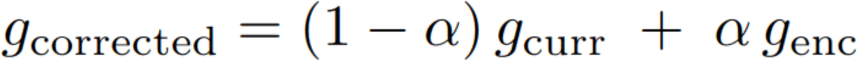

Here, g_corrected_ is the grid cell population vector after correction, g_curr_ is a noisy current grid cell population vector and g_enc_ is the grid cell population vector stored during encoding. α controls the level of correction. If the distance between the encoded and current positions is larger than 0.5 meters, no correction was applied.

In the incongruent case, we estimated the mean and standard deviation of normal distributions for the x and y step components by fitting them to experimental data from the incongruent phase. Instead of fitting distributions for each step length and leg side, we fitted them for each trajectory distance (1.2 m, 2.2 m, 3.4 m). Because the incongruent condition involves more regular step patterns that do not match any stored tuples, we could not retrieve a concatenated vector from memory. No error-correction was applied in this scenario.

The model was implemented in Python (version 3.11.4) with Numpy, Matplotlib, and Pandas libraries.

## Acknowledgments

We thank Max Schulz and Paula Schneider for helping with the data collection and Kiran Varanasi for technical advice. We also thank Misun Kim and colleagues of the Department of Psychology at MPI CBS, Leipzig, for prolific discussions. This work was supported by the Max Planck Society, the Kavli Foundation, the Jebsen Foundation and Helse Midt Norge, awarded by C.F.D.

## Author contribution

*VR:* Project idea, Conceptualization, Task programming, Data acquisition, Formal analysis, Visualization, Software, Writing - original draft, Writing - review & editing

*LK:* Conceptualization, Task programming, Formal analysis, Visualization, Software, Writing - review & editing

*VS:* Modeling, Visualization, Writing - original draft, Writing - review & editing

*MB:* Conceptualization, Supervision, Writing - review & editing

AB: Modeling, Supervision, Writing - review & editing

*CD:* Conceptualization, Supervision, Resources, Writing - review & editing

## Data availability

Data to reproduce the statistical analyses reported in this paper will be made available upon publication via the Open Science Framework.

## Code availability

The analysis code will be available upon publication on GitHub.

## Supplementary materials

**Figure S1.**
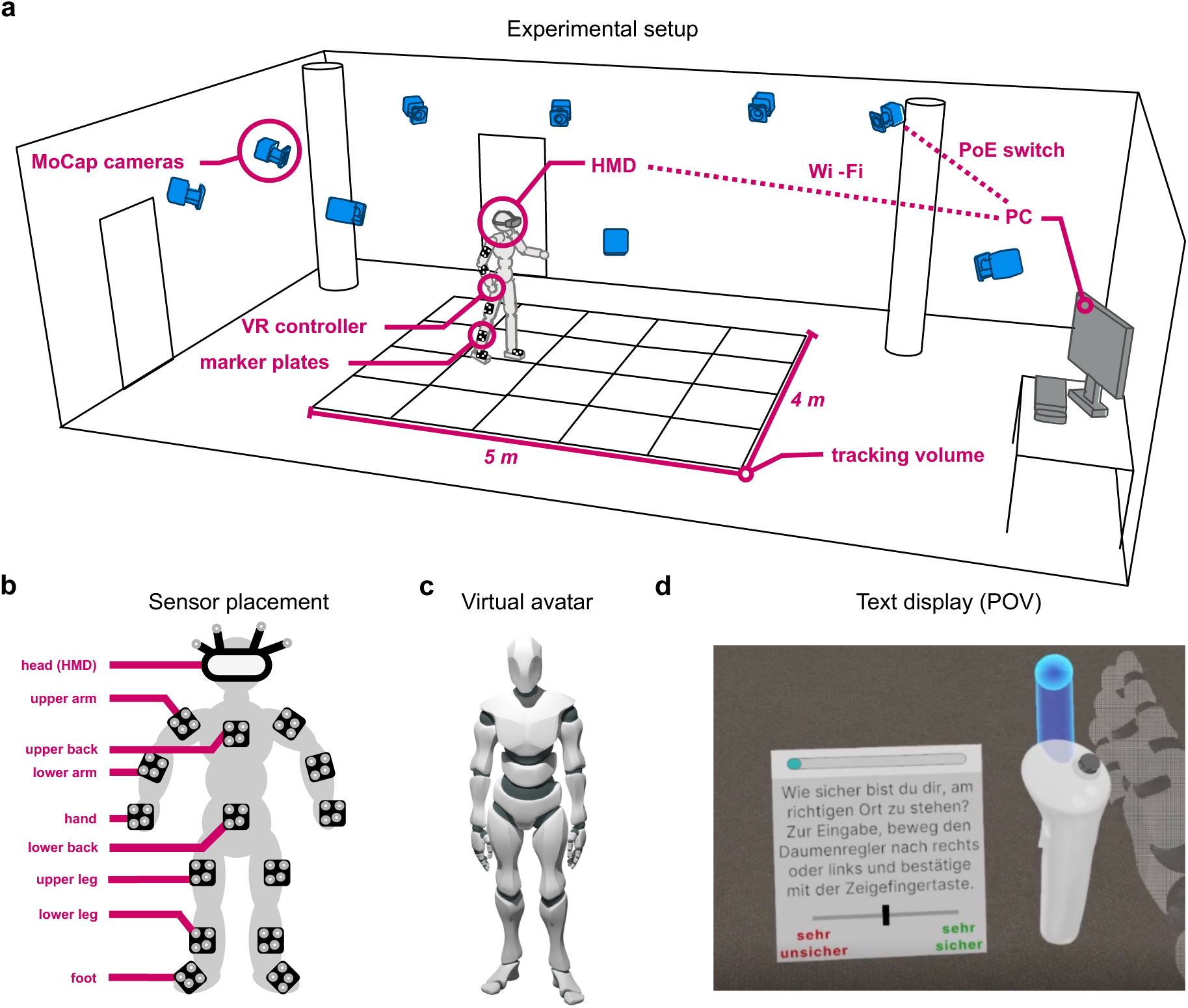
Experimental setup. **a:** Illustration of the MoCap-integrated VR laboratory setup and the equipment used. **b:** Placement of rigid body markers on various parts of the body. c: Animated humanoid avatar that was mapped onto each participant’s body with individual scaling. d: First-person view of the virtual text display used to present task instructions and collect confidence ratings.

**Figure S2.**
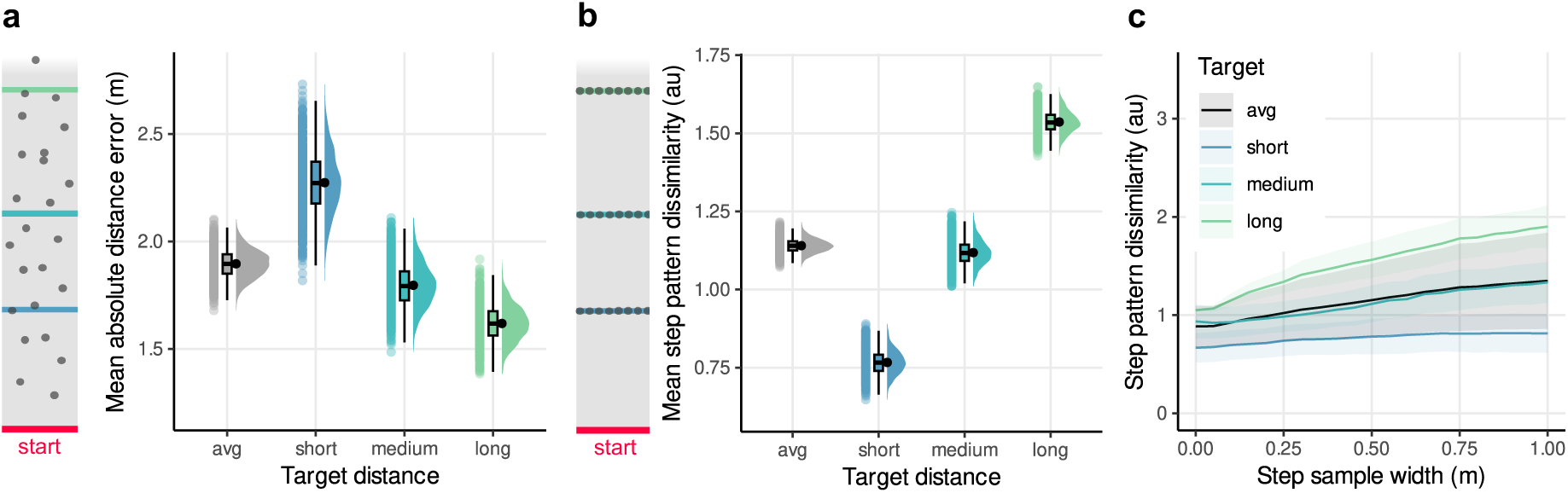
Simulation-based generation of chance levels. **a:** Absolute distance errors. *Left:* Random responses (gray dots) for each target distance (colored lines) along the linear track were sampled from a uniform distribution, spanning 0.05 to 6.5 m from the start (red line). *Right:* Based on these responses, we calculated absolute distance errors for 1,000 artificial participants, each following the original trial structure of 6 trials per, resulting in a chance distribution for each target distance and the average. Violin plots depict the density distribution, boxplots the median and quartiles, black dots with error bars the means ± SEM, and colored dots individual artificial participants per condition. **b:** Step pattern dissimilarity (SPD). *Left:* Random step patterns were sampled between the start and target locations, preserving the same number of steps as during encoding. *Right:* Accordingly, we calculated SPD values for 1,000 artificial participants, each following the original trial structure of 6 trials per, resulting in a chance distribution for each target distance and the average. **c:** SPD as a function of step sample width for each target distance and the average. The step sample width indicates the regularity by which steps can be performed to reach a given target. A step sample width of 0 reflects perfectly equal sized steps (target distance / correct number of steps). With a step sample width of 1, steps were uniformly sampled in an area of 1 meter around the regular step length. Across the 1,000 artificial participants, the step sample width was divided into 10 bins ranging from 0 to 1 meter. Colored lines reflect the mean ± SD (ribbon) per condition.

**Figure S3.**
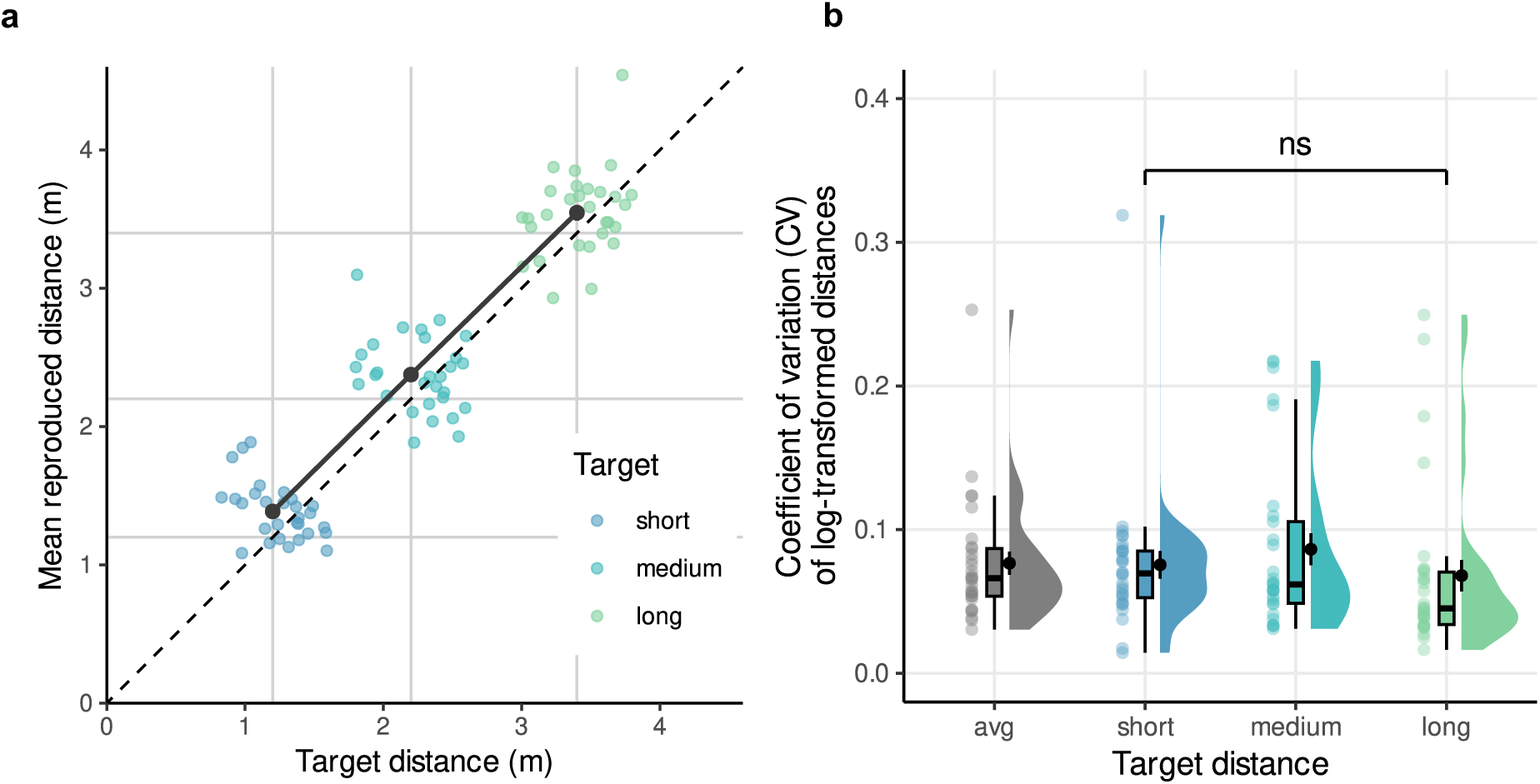
Reproduced versus target distance and precision. **a:** Mean reproduced distance as a linear function of target distance. A bootstrapped 1-way repeated measures ANOVA revealed a significant effect of target distance on reproduced distance (*F*1.91,53.5 = 1148, *P* < 0.0001, η^2^_*p*_= 0.98, 95% CI = 0.97 to 0.99) with all pairwise comparisons being significant (bootstrapped paired *t*-tests: all *Padj*’s < 0.003, Bonferroni-corrected for 3 comparisons). Black dots with error bars depict the means ± SEM, and colored dots individual participants per condition. **b.** Coefficient of variation (CV) as function target distance. A bootstrapped 1-way repeated measures ANOVA showed no significant effect of target distance on log-transformed CV (*F*1.9,52.8 = 1.22, *P* = 0.31, η^2^_*p*_= 0.04, 95% CI = 0.003 to 0.23). Violin plots depict the density distribution, and boxplots the median and quartiles.

**Figure S4.**
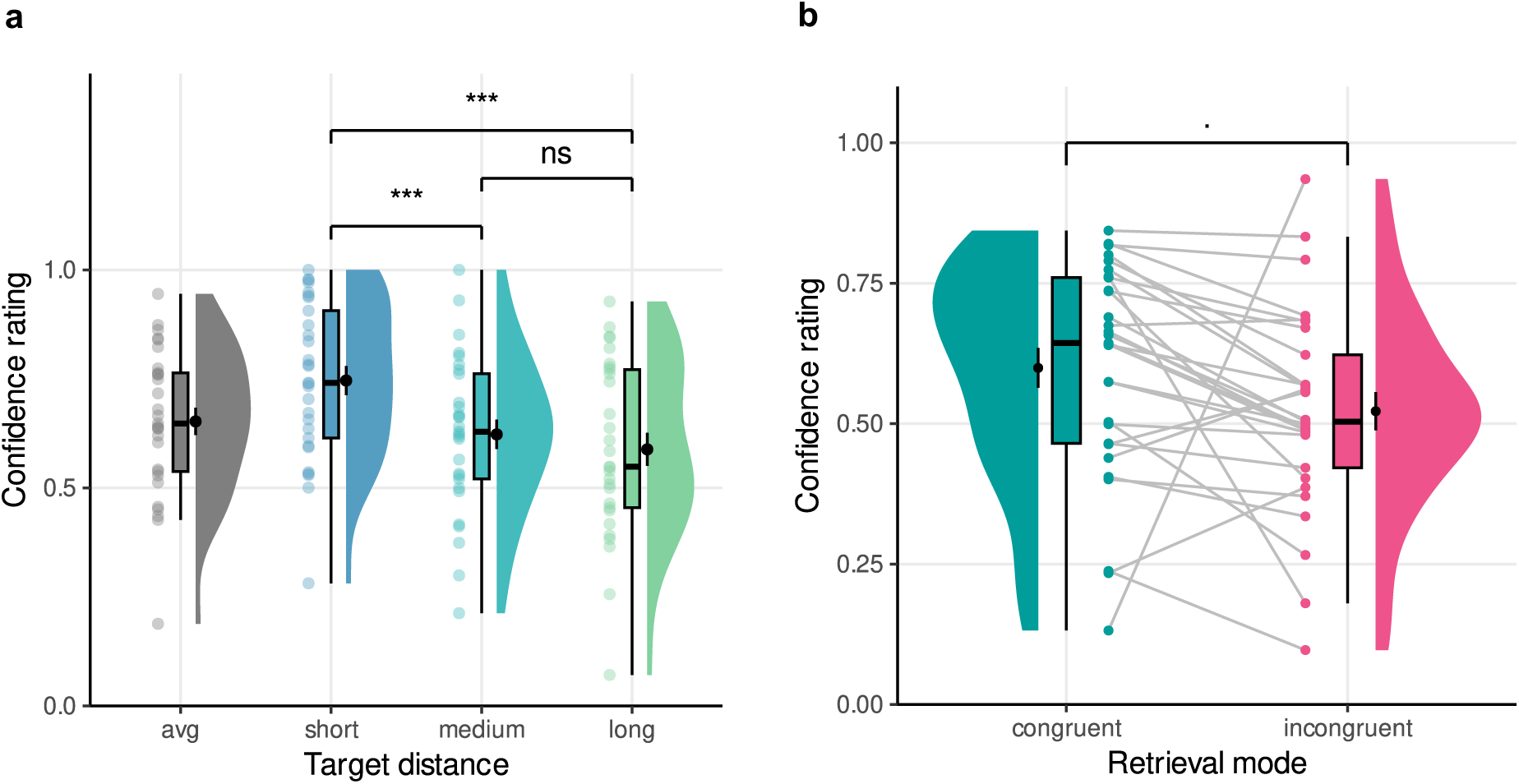
Confidence ratings. **a.** Confidence ratings (ranging from “very uncertain” = 0 to “very certain” = 1) as a function of target distance during free retrieval. A bootstrapped 1-way repeated measures ANOVA revealed a significant effect of target distance on confidence ratings (*F*1.91,53.56 = 18.81, *P* < 0.0001, η^2^_*p*_ = 0.40, 95% CI = 0.28 to 0.55). Post-hoc bootstrapped paired *t*-tests indicate pairwise differences between short and long as well as short and medium (all *P*’s < 0.0001). Violin plots depict the density distribution, boxplots the median and quartiles, black dots with error bars the means ± SEM, and colored dots individual participants per condition. **b.** Confidence ratings as a function of retrieval mode. A bootstrapped paired *t*-test revealed a non-significant statistical trend for higher response confidence during congruent vs incongruent retrieval (*t*28 = 1.86, *P* = 0.09, *d* = 0.35, 95% CI = -0.02 to 1.14). ᐧ *P* < .01, *** *P* < .001.

**Table S1.** Sample characteristics

| **Variable** | **Frequency** | **Percentage (%)** |
| --- | --- | --- |
| **Gender** |  |  |
| Female | 12 | 41.38 |
| Male | 14 | 48.28 |
| Non-binary | 3 | 10.34 |
| **Age (years)** |  |  |
| 18-23 | 7 | 24.14 |
| 24-29 | 13 | 44.83 |
| 30-35 | 9 | 31.03 |
| **Education** |  |  |
| < High school diploma | 0 | 0.00 |
| High school diploma | 12 | 41.38 |
| Bachelor's degree | 11 | 37.93 |
| Master's degree | 4 | 13.79 |
| > Higher univ. degree | 0 | 0.00 |

**Table S2.** Body measures.

| **Variable** | **Mean ± SD (min-max)** |
| --- | --- |
| Weight (kg) | 70.552 ± 13.571 (50.0 - 99.0) |
| Height (m) | 1.746 ± 0.067 (1.64 - 1.86) |
| Arm span (m) | 1.726 ± 0.0743 (1.61 - 1.86) |
| Inseam (m) | 0.783 ± 0.033 (0.73 - 0.86) |
| Hip width (m) | 0.386 ± 0.093 (0.29 - 0.85) |

**Table S3.**
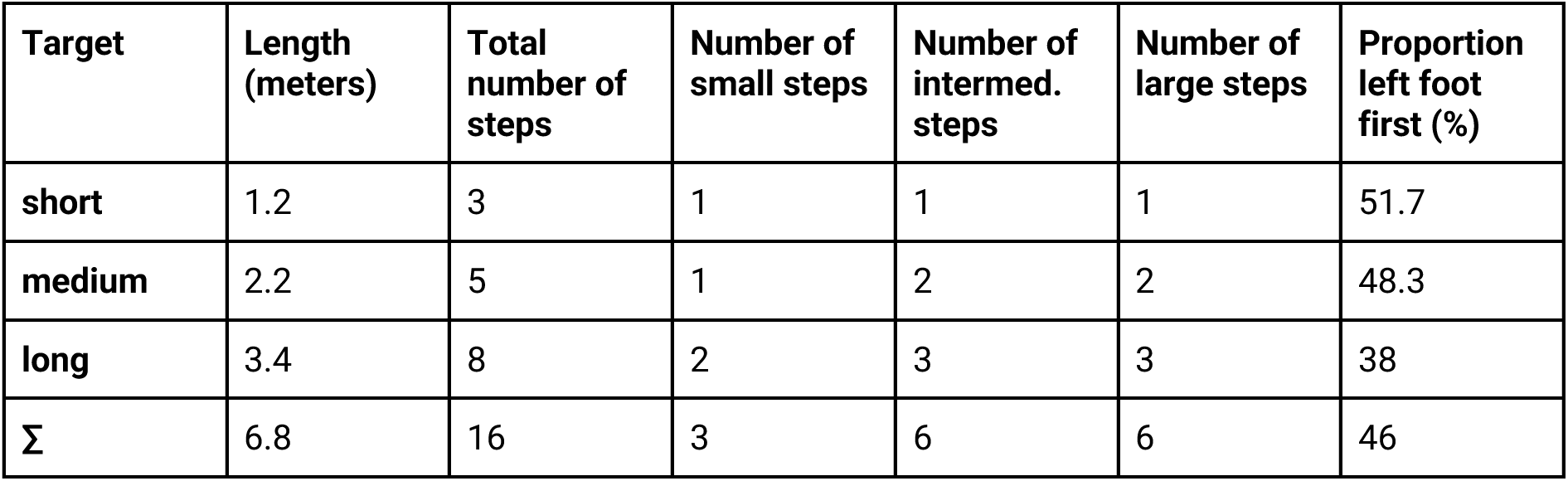
Encoding step pattern characteristics.

| **Target** | **Length**<br>**(meters)** | **Total**<br>**number of**<br>**steps** | **Number of**<br>**small steps** | **Number of**<br>**intermed.**<br>**steps** | **Number of**<br>**large steps** | **Proportion**<br>**left foot**<br>**first (%)** |
| --- | --- | --- | --- | --- | --- | --- |
| **short** | 1.2 | 3 | 1 | 1 | 1 | 51.7 |
| **medium** | 2.2 | 5 | 1 | 2 | 2 | 48.3 |
| **long** | 3.4 | 8 | 2 | 3 | 3 | 38 |
| **Σ** | 6.8 | 16 | 3 | 6 | 6 | 46 |

**Table S4.** Debriefing summary

| **Item** | **M ± SD \| n/N (%)** |
| --- | --- |
| **A: Target distance (au)** |  |
| *How long was the short target distance?* | 0.15 ± 10.08 |
| *How long was the medium target distance?* | 0.29 ± 0.14 |
| *How long was the long target distance?* | 0.5 ± 0.24 |
| *How many step lengths were presented?* | 3.43 ± 0.6 |
| **B: Number of steps (frequency)** |  |
| *How many steps occurred in the short target distance?* | 3.83 ± 0.57 |
| *How many steps occurred in the medium target distance?* | 5.83 ± 0.47 |
| *How many steps occurred in the long target distance?* | 8.76 ± 0.58 |
| **C: Strategies (frequency)** |  |
| *I imagined the target positions relative to the start* | 8/29 (27.59%) |
| *I imagined the target positions relative to each other* | 16/29 (55.17%) |
| *I counted the seconds until I reached my destination* | 0/29 (0%) |
| *I remembered my body movements* | 25/29 (86.21%) |
| *I used ambient noise for orientation in VR* | 2/29 (6.90%)<sup>a</sup> |
| *I orientated myself based on the physical room layout* | 0/29 (0%) |
| **D: Body attention (frequency)** |  |
| *I paid the most attention to the head* | 11/29 (37.93%) |
| *I paid the most attention to the torso* | 2/29 (6.90%) |
| *I paid the most attention to the upper arms* | 1/29 (3.45%) |
| *I paid the most attention to the lower arms* | 1/29 (3.45%) |
| *I paid the most attention to the hands* | 2/29 (6.90%) |
| *I paid the most attention to the hips* | 4/29 (13.79%) |
| *I paid the most attention to the upper legs* | 15/29 (51.72%) |
| *I paid the most attention to the lower legs* | 13/29 (44.83%) |
| *I paid the most attention to the feet* | 27/29 (93.10%) |
| **E: Volition (au)** |  |
| *How consciously did you try to repeat the original step patterns?* | 0.77 ± 0.29 |
| **F: Balance (au)** |  |
| *How difficult was it for you to maintain the balance during encoding?* | 0.6 ± 0.22 |
| *How difficult was it for you to maintain the balance during retrieval?* | 0.6 ± 0.20 |
| **G: Motion sickness (au)** |  |
| *To what extent did you experience symptoms such as dizziness, headaches or nausea?* | 0.2 ± 0.26 |
| **H: Motivation (au)** |  |
| *How highly do you rate your motivation during the task?* | 0.8 ± 0.16 |

Note. Response formats. A: Drag and drop tags (color-coded target locations) onto a virtual track spanning from the left (min = 0) to the right (max = 1) screen side from birds-eye-view; B: Digit input box (min = 0; max = inf); C: Checkboxes (multiple choice); D: Checkboxes (multiple choice); E: Continuous slider ranging from "very unconscious" (min = 0) to "very conscious" (max = 1); F: Continuous slider ranging from "very easy" (min = 0) to "very hard" (max = 1); G: Continuous slider ranging from "not at all" (min = 0) to "very strong" (max = 1); H: Continuous slider ranging from "very low" (min = 0) to "very much" (max = 1).
<sup>a</sup> When asked, both participants added that auditory information did *not* help them to estimate distance.

